# *Dlx5/6* regulate perineuronal net-synapse coupling and stabilize adult cortical Parvalbumin neurons networks

**DOI:** 10.64898/2026.07.23.740286

**Authors:** Lou Belz, Rym Aouci, Baptiste Allain, Fiona Henderson, Anastasia Fontaine, Pénélope Desgagné, Laure Calzaroni, Véronique Fabre, M. Belén Pardi, Giovanni Levi, Nicolas Narboux-Nême

**Affiliations:** Molecular Physiology and Adaptation, National Museum of Natural History (MNHN), CNRS UMR7221, 75005, Paris, France; Université Paris Cité, Institute of Psychiatry and Neuroscience of Paris (IPNP), INSERM U1266, Neuronal circuits for memory and perception Team, 75014, Paris, France; Sorbonne Université, INSERM, CNRS, Institut de Biologie Paris Seine (IBPS), Center for Neuroscience at Sorbonne Université (NeuroSU), 75005, Paris, France

## Abstract

The transcriptional mechanisms that maintain adult Parvalbumin (PV) interneuron function and cortical network stability remain poorly understood. We previously showed that the inactivation in GABAergic neurons of *Dlx5/6*, coding for transcription factors, disrupts social interaction and reduces PV interneuron density in the prefrontal cortex. Here, combining transcriptional and histological analyses with *ex vivo* electrophysiological and *in vivo* electroencephalographical (EEG) recordings, we show that *Dlx5/6* regulate a molecular program controlling perineuronal net (PNN) homeostasis in the adult cortex. *Dlx5/6* inactivation dysregulated the expression of multiple PNN-associated genes and induced region-specific remodelling of PNN mesh architecture in the prefrontal and somatosensory cortices. These structural changes were accompanied by alterations in excitatory and inhibitory synaptic organization and by a disrupted coupling between local PNN structure and synaptic properties. *Ex vivo* electrophysiological recordings revealed that fast-spiking interneurons displayed altered intrinsic properties and reduced excitatory synaptic drive in a region-specific manner, while EEG recordings during social interaction showed impaired recruitment of prefrontal gamma oscillations. Together, these findings identify *Dlx5/6* as regulators of adult PV interneuron stability, linking extracellular matrix homeostasis to synaptic organization and cortical network dynamics. More broadly, it provides a new mechanistic framework connecting *Dlx5/6* function to PV-related pathological phenotypes, including neuropsychiatric disorders.

**GRAPHICAL ABSTRACT:** 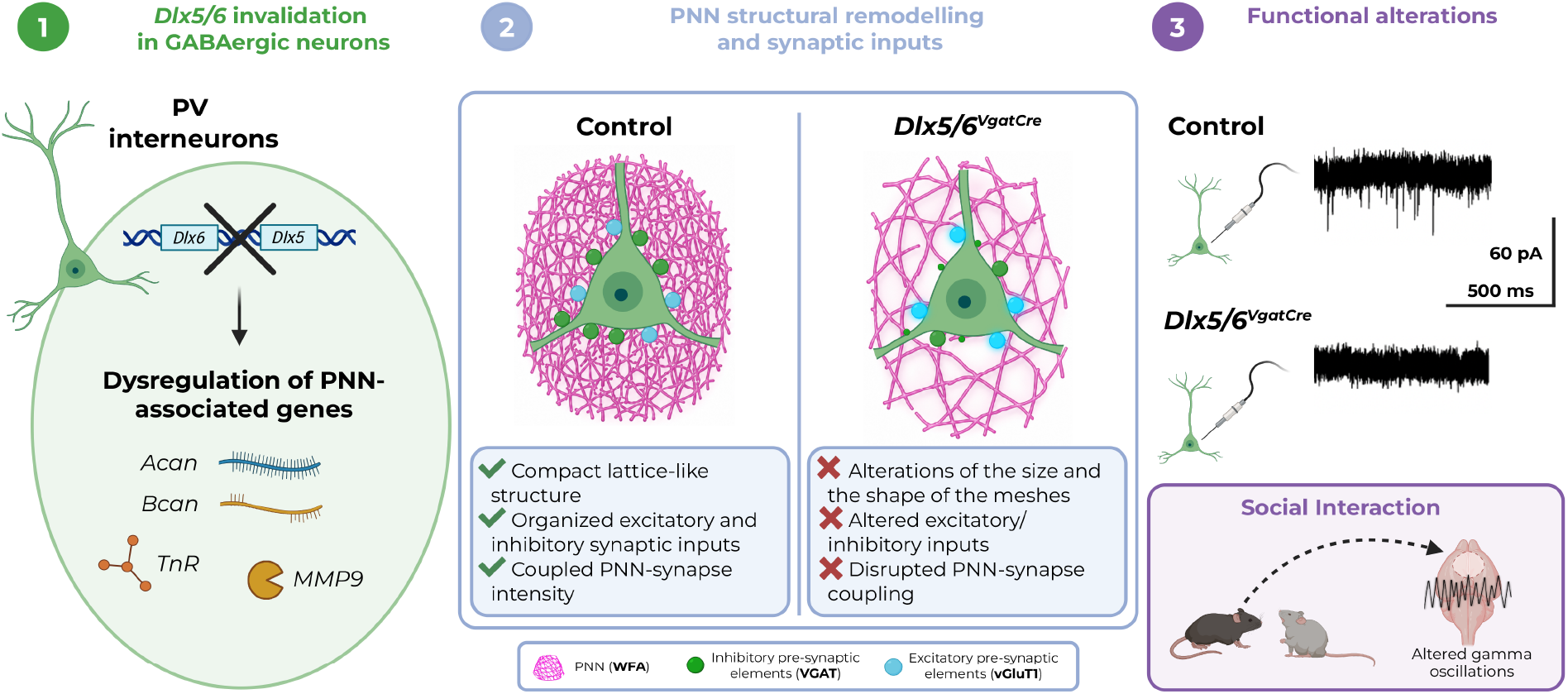

## INTRODUCTION

Cortical inhibitory circuits are essential for the generation of higher-order brain functions, including perception, cognition and social behaviour^1–3^. Among GABAergic interneurons, Parvalbumin (PV)-expressing neurons play a central role in coordinating cortical network activity and supporting complex behaviours^4,5^. Yet the cellular and molecular mechanisms that stabilize these networks and allow their adaptive modulation remain poorly understood. Our previous studies on the homeobox genes *Dlx5* and *Dlx6* established the role of transcriptional programs in cortical PV neuronal density and behaviours^6–8^.

While *Dlx5/6* are well established as key developmental transcription factors^9,10^, their role in maintaining mature inhibitory circuits after cortical maturation remains largely unexplored. Whether *Dlx5/6* participate in transcriptional programs that preserve PV interneuron stability and cortical network function throughout adulthood remains to be fully established. Initial evidence for the importance of *Dlx5/6* in PV interneuron differentiation was provided by heterochronic grafting experiments showing that *Dlx5/6*-null GABAergic neuroblasts failed to efficiently differentiate into PV-positive interneurons after transplantation into the postnatal cortex^11^. Consistent with this observation, constitutive *Dlx5/6* heterozygous mice display abnormalities in prefrontal gamma oscillations, a network activity critically dependent on PV interneuron synchrony^12^. However, because *Dlx5/6* are also required for the development of several non-neural organs, constitutive deletion of these genes is embryonic lethal, precluding the study of their function in the adult nervous system. To overcome this, we selectively manipulated *Dlx5/6* dosage in GABAergic neurons, generating mouse models that display robust alterations in social interaction, vocal communication and anxiety-related behaviours^6–8^, suggesting an important participation of *Dlx5/6* in the organization of neuronal networks underlying complex behaviours, including social behaviour.

This genetic strategy provided a powerful framework to investigate the contribution of these genes to the organization and function of adult inhibitory networks. We previously reported a linear correlation between *Dlx5/6* allelic dosage and the density of PV neurons in the adult prefrontal cortex, also linearly associated with anxiety- and compulsive-like behaviours^8^.

PV interneurons, including basket and chandelier cells, provide powerful perisomatic inhibition and are key determinants of cortical network synchrony^13,14^. Their characteristic fast-spiking electrophysiological properties underlie the coordinated firing that gives rise to high-frequency gamma oscillations across cortical networks^15,16^. Alterations in PV interneuron function and gamma oscillatory activity have been consistently associated with neuropsychiatric disorders, including autism spectrum disorders and schizophrenia^17,18^.

The functional stability of PV interneurons in the adult cortex depends not only on the expression of specific ion channels^19^ and membrane proteins, but also on a specialized extracellular matrix structure known as the perineuronal net (PNN), which predominantly enwraps the soma and proximal dendrites of PV interneurons. This lattice-like structure contributes to the stabilization of cortical circuits by regulating synaptic organization, neuronal excitability and the electrophysiological properties of PV interneurons^20,21^.

PNNs serve multiple functions: they constrain synaptic inputs, regulate the perisomatic ionic homeostasis and protect neurons from oxidative stress^20,22,23^. PNNs participate in synapse maintenance by interacting with pre- and post-synaptic elements as well as surrounding astrocytes^24^; they are considered by some as integral components of the “tetrapartite synapse” complex^25^. Their core scaffold consists of hyaluronic acid (HA) chains synthesized by Hyaluronan Synthases (HASs), anchored to the cell surface. Chondroitin sulphate proteoglycans (CSPGs) are attached to this scaffold through hyaluronan and proteoglycan link proteins (HAPLNs). The CSPGs composition varies according to the lectican family member involved: Brevican (Bcan), Neurocan (Ncan), Versican (Vcan), and Aggrecan (Acan), the latter generally most abundant^26,27^. Structural rigidity is conferred by crosslinking proteins, most prominently Tenascin-R (TnR)^26^. PV interneurons themselves contribute to PNN component synthesis, with additional production from astrocytes and oligodendrocytes of assembly and remodelling enzymes^26^.

PNNs are highly dynamic structures: HAS continuously synthesizes new hyaluronic acid scaffold, while extracellular proteases, including Matrix metalloproteinase (MMPs) and ADAMTSs (a disintegrin and metalloprotease with thrombospondin motifs), mediate ongoing CSPGs cleavage and remodelling^28,23^.

Beyond its recognized role in PV interneuron stability and cortical plasticity, PNNs dysfunction has been recently involved in the pathophysiology of several neuropsychiatric disorders, including schizophrenia, bipolar disorder, and autism spectrum disorder^29^. Yet, the transcriptional mechanisms that maintain PV properties and PNN homeostasis in the adult brain remain poorly understood.

In the present study, we used a conditional mouse model in which *Dlx5/6* is selectively inactivated in GABAergic neurons (*Dlx5/6^VgatCre^*) to investigate how these transcription factors contribute to the maintenance of PV interneuron properties and cortical network dynamics in the adult brain. Combining molecular assays, quantitative analyses of PNNs architecture, synaptic profiling, *ex vivo* electrophysiology and EEG recordings during social behavioural engagement, a task known to enhance gamma oscillations, we show that *Dlx5/6* regulate a transcriptional program linking extracellular matrix homeostasis to synaptic organization and inhibitory network synchronization. We performed analyses in two cortical regions with distinct functional profiles: the somatosensory barrel cortex (S1BF), a primary sensory area with high PV interneurons and PNN densities^30^, and the prelimbic cortex (PrL), a prefrontal region in which we previously shown that *Dlx5/6* inactivation PV interneurons density^8^. Our findings show that a *Dlx5/6* deficit modifies adult PNNs architecture and PV interneuron stability, affecting the dynamic recruitment of gamma oscillations that underlies adaptive social behaviour.

## MATERIALS AND METHODS

### Animals

All animal procedures were conducted in accordance with European and French Agriculture Ministry directives. Experimental protocols were reviewed and approved by the MNHN “Cuvier” ethics committee and validated by the “Ministère de l’Enseignement Supérieur et de la Recherche” (Apafis #1612).

Mice were housed under controlled light with a 12 h light/dark cycle, constant temperature (21 °C), and humidity (50–60%). Food and water were available *ad libitum*.

*Dlx5/6^lox/lox^* mice were crossed with *Vgat^cre/+^* mice to generate *Dlx5/6^VgatCre/+^*and then *Dlx5/6^VgatCre^*, as previously described^7^. This strategy enabled selective inactivation of *Dlx5/6* in GABAergic neuroblasts. Since in the telencephalon, *Dlx5/6* is only expressed in ganglionic eminences derived GABAergic neurons, the cortical phenotype results from GABAergic neurons *Dlx5/6* inactivation.

In all experiments, *Dlx5/6^lox/lox^* littermates with no *Vgat^cre^* allele were used as controls of *Dlx5/6^VgatCre^*mice. The *Vgat^cre^* mouse line is a *Vgat-Ires-Cre* knock-in, and previous RT-qPCR experiments have verified that *Vgat* expression is maintained in these mice.

### Reverse transcription quantitative PCR (RT-qPCR)

Cortical fragments from the somatosensory and prefrontal areas were microdissected from 0.5-1 mm-thick brain sections under a stereomicroscope in cold Tris-HCl (0.1 M, pH 7.6) buffer and immediately frozen on dry ice. Tissue fragments were homogenized using a tissue-Lyser system (QIAGEN, France) and total RNA was isolated using a RNAquerous-Micro Kit (Invitrogen, ref: 1931).

To eliminate potential DNA contamination, samples were treated with DNase using the Kit Turbo DNA free (Ambion, Appliedbiosystems, ref: AM1907). RNA quality was assessed using Agilent Microfluidic chips with the RNA 6000 Pico Kit for low concentration samples (Agilent, ref: 5067-1513).

cDNA was synthesized from 1 µg of RNA using reverse transcriptase reagents (Invitrogen, ref: 10777-019). Real-time quantitative PCR (RT-qPCR) was performed using the SYBR Green method (*Power* SYBR^TM^ Green PCR Master Mix, ref.: 4367659 Applied Biosystems).

Relative gene expression was calculated using the comparative Ct (2-ΔΔCt) method. Normalized expression values relative to the reference genes *β-actin* and *β3-tubulin* were determined using MxPro qPCR software (Agilent Technologies).

Primers sequences are listed in Supplementary Table 1.

### Immunohistochemistry (IHC)

Animals were euthanized with an overdose of pentobarbital injection (1.5g/kg Pentobarbital, IP, Euthasol Dechra). After respiratory arrest, they were intracardially perfused with 4% paraformaldehyde in phosphate buffer. Brains were removed, postfixed overnight at 4 °C in the same fixative, cryoprotected by immersion in 30% sucrose for 48 h incubation at 4°C, frozen and stored at −20°C until sectioning. Free-floating coronal section (60 µm thick) were using a Leica cryostat (CM3050S). Sections were either processed immediately for immunohistochemistry or stored at −20°C in a cryoprotective solution (30% Glycerol, 30% Ethylene glycol, PBS1X).

Brain sections were incubated for 48 h incubation at 4 °C with primary antibodies diluted in blocking solution (PBS 1%, 2% gelatine and 0.25% Triton); anti-Parvalbumin (1:2000, Invitrogen, ref.: Pa1-933); anti-VGLT1 (1:2000, Synaptic System, ref.: 135 303); anti-VGAT (1:200, Synaptic System, ref.: 131 011); Lectin from Wisteria floribunda (1:500, Sigma, L1516-2MG). Perineuronal Net (PNNs) were visualized using biotin-linked Wisteria floribunda agglutinin (WFA) labelling, which links to the chondroitin sulphate-rich structures of the extracellular matrix^31^.

Species matching Alexa 488 and 594 fluor-conjugated F(ab’)2 secondary antibodies (1:500 Jackson Immunoresearch) were used to reveal primary antibodies CY3-conjugated and Alexa Fluor 647-conjugated Streptavidin (Jackson Immunoresearch) were used to reveal biotin-WFA. Sections were incubated in the presence of DAPI (1:1000) to label nuclei Sections were mounted using Fluoromount-G mounting medium (ThermoFisher, France).

### Acquisitions and density analysis of PV^+^PNN^+^ neurons

Epifluorescence imaging of brain sections was performed with a Zeiss Macro-Apotome Axiozoom V16, equipped with the Apotome Module. Cortical regions and cortical layers were identified using DAPI counterstaining according to the Paxinos and Franklin mouse brain atlas^32^.

PV-positive and WFA-positive neurons were manually counted using the Fiji ImageJ cell counter plugin. Cell density was calculated by normalizing the number of labelled cells to the analysed surface. Quantification was performed bilaterally and averaged across hemispheres. All analyses were performed blind to genotype. The number of cells and animals analysed are indicated for each analysis.

### Acquisitions and analysis of the Perineuronal Net structure and synaptic contacts onto PV somata

Images of Parvalbumin PV^+^ neurons and Wisteria floribunda agglutinin WFA^+^ neurons were acquired using a Zeiss LSM 880 confocal microscope equipped with Airyscan detector (Carl Zeiss). High-resolution imaging of PNNs was performed using a 63× oil-immersion objective (NA 1.4). Z-stacks were acquired with a step size of 0.44 µm.

All analysed PNNs were located within layer V of the somatosensory and prefrontal cortices. Cortical regions and layers were identified according to the Paxinos and Franklin mouse brain atlas^32^. Imaging parameters, including laser power, detector gain, pinhole diameter, and acquisition settings, were kept constant across all imaging sessions and genotypes.

Image processing and quantitative analyses were performed using the APNU2.0 macro implemented in ImageJ/Fiji software, by Dr Dityatev’s team^33^. Quantifications were performed independently on the left and right hemispheres and subsequently averaged for each animal.

To assess synaptic organization around PNNs, we used a semi-automatic Fiji-based analysis pipeline derived from APNU2.0.^34^ As the original APNU2.0 script is designed to analyse single-channel images, we implemented an additional plugin, provided by Dr Dityatev, enabling the simultaneous analysis of two channels. This modification enabled the selective quantification of synaptic puncta ensheathed by PNNs.

Images were acquired in three channels: Wisteria floribunda agglutinin (WFA) labelling PNNs, VGLUT1 immunostaining for excitatory presynaptic terminals, and VGAT immunostaining for inhibitory presynaptic terminals. For quantitative analysis, images were processed as paired-channels combinations (WFA^+^ VGLUT1 or WFA^+^ VGAT). This approach enabled comparative quantifications of excitatory and inhibitory synaptic puncta associated with the same PNN structures.

### Surgery and *in vivo* Electrode Implantations

Mice were anesthetized with a mix of ketamine/xylazine (100 and 10 mg/kg, respectively, IP) and positioned in a stereotaxic frame. Stereotaxic coordinates were determined according to Paxinos and Franklin mouse brain atlas^32^.

Mice were implanted with electrodes fabricated from enameled nichrome wire (150 µm in diameter) to enable chronic electroencephalographic (EEG) and electromyographic (EMG) recordings. Four EEG electrodes were placed epidurally via small burr holes drilled through the skull at the following coordinates: right frontal cortex (2 mm lateral to midline, 2 mm anterior to bregma), right parietal cortex (2 mm lateral to midline, 2 mm posterior to bregma), left parietal cortex (2 mm lateral to midline, 2 mm posterior to bregma), and cerebellum (midline, 2 mm posterior to lambda). Two additional EMG electrodes were bilaterally inserted into the nuchal muscles. This electrode configuration yielded fronto-parietal EEG and parietal-cerebellar EEG derivations, together with an EMG derivation. All electrodes were soldered to a miniature connector (Antelec, La Queue-en-Brie, France) and secured to the skull with dental acrylic cement. Following surgery, mice were allowed a 10-day recovery period and were handled daily to facilitate habituation. Prior to EEG/EMG recordings, mice were acclimated for 4 days to the recording environment in individual Plexiglas chambers (19 × 19 × 30 cm). During both acclimation and recording sessions, animals were connected to the acquisition system via a lightweight cable suspended from a swivel commutator, permitting unrestricted movement within the chamber.

### Social Interaction Task and recordings

A juvenile conspecific (3-4 weeks old, same sex as the experimental animal) was introduced into the recording chamber of the test mouse. The two freely moving animals were allowed to interact for 5 min. EEG/EMG signals were continuously acquired using an Embla amplifier (Medcare, Reykjavik, Iceland), and animal behaviour was simultaneously video-monitored through the same system.

### Scoring

EEG and EMG signals were amplified (2,000×), bandpass filtered, digitized at 2,000 Hz, and subsequently downsampled to 200 Hz. Signals and video recordings were scored offline to identify and quantify social behaviours directed toward the juvenile intruder. Social interactions, including sniffing, close following and allogrooming, were scored in 3 s epoch using Somnologica®software (Medcare, Reykjavik, Iceland).

### EEG analysis

EEG signals were processed for Power spectral density (PSD) analysis using the Animal Spectrogram function in Somnologica®software (Medcare, Reykjavik, Iceland). Epochs containing artifacts were excluded from the analysis. Spectral estimates were computed using a fast Fourier transform (FFT) with a window size of 1024 points, yielding a frequency resolution of 0.2 Hz and power estimates expressed in µV²/Hz. The 48-52 Hz frequency range was excluded from all analyses to avoid contamination by electrical line noise artefacts. Analyses were performed for both the social interaction condition and baseline periods, the latter corresponding to epochs without active exploration. To characterize baseline spectral profiles and compare relative power across genotypes and frequency bands, PSD values for each animal were normalized to the total spectral power within the 0.5-80 Hz range (excluding 48-52 Hz) and expressed as relative power (%). Group spectra were generated by averaging normalized spectra within each genotype group (Ctrl and *Dlx5/6^VgatCre^*) and are presented as mean ± SEM of log_10_-transformed relative power values.

To assess task-evoked changes in oscillatory activity, unnormalized power during the social interaction task was compared with baseline power for each animal. At each frequency bin (*f*_i_), power modulation was calculated as: Ratio(fᵢ) = 10 × log_10_ (PSD_task(f_i_) / PSD_baseline(f_i_)) and expressed in decibels (dB), where positive values indicate increased power during the task relative to baseline, and negative values indicate decreased power. For band-specific analyses, mean ratio values were calculated by averaging all frequency bins within each frequency band for each individual animal: delta (0.5–4.99 Hz), theta (5–9.99 Hz), alpha (10–12.99 Hz), beta (13–29.99 Hz), low gamma (30–48 Hz), and high gamma (52–80 Hz).

### *In vitro* electrophysiology Slice preparation

To enable the recording of PV-positive neurons, mice were injected intravenously with an AAV vector (AAV-PHP.eB/2-S5E2-HBB-chl-dTomato_2A_NLSdTomato), which selectively allows the expression of dTomato in fast-spiking interneurons. Briefly, 2.5×10^10^ viral particles of Brain-blood barrier crossing AAV were injected in the tail vein 3 weeks prior to the experiment. dTomato expression was under the control of the S5E2 enhancer, enabling selective labelling of Nav1.1-positive fast-spiking interneurons^35^. The specificity of the S5E2 enhancer for cortical PV/fast-spiking interneurons had previously been validated^35^.

Brain slice preparation was adapted from a protocol optimized for adult mice3. Following cervical dislocation, brains were rapidly removed and immersed in ice-cold N-Methyl-D-Glutamine solution (NMDG)-based cutting solution containing (in mM): 93 NMDG, 2.5 KCl, 1.2 NaH_2_PO_4_, 30 NaHCO_3_, 20 HEPES, 25 Glucose, 5 Na ascorbate, 2 Thiourea, 3 Na pyruvate, 10 MgSO_4_, 0.5 CaCl_2_, pH = [7.3;7.4]). Brains were maintained in ice-cold NMDG solution throughout extraction and slicing procedures. Following sectioning, slices were incubated in warm NMDG solution (34°C) for 9-13 minutes and subsequently transferred to artificial cerebrospinal fluid (ACSF) containing (in mM): 125 NaCl, 3 KCl, 1.25 NaH_2_PO_4_, 10 Glucose, 26 NaHCO_3_, 1 MgSO_4_, 2 CaCl_2_, continuously oxygenated with 95% O_2_ – 5% CO_2_. Slices were maintained in ACSF at room temperature for at least 30 min before recording. Recordings were performed in both hemispheres. Slices were transferred to the recording chamber and neurons were visualized using differential interference contract (DIC) microscopy on a SliceScope Pro 2000 microscope (Scientifica^TM^) equipped with a 40X water immersion objective (Olympus^TM^) and a charge-coupled device (CCD) camera (Scientifica^TM^/QI Imaging Ocular^TM^). Fluorescent cells were identified using LEDs illumination (CoolLED^TM^ pE-300 ultra) through the objective.

### Intracellular recording

Whole-cell recordings were performed at room temperature. Signals were amplified through Axon^TM^ Digidata 1550B amplifier and then digitized at 20 kHz and low-pass filtered at 10 kHz by a MultiClamp^TM^ 700B digital-to-analog converted. Recordings were excluded from analysis when access resistance exceeded 30 MΩ. Membrane potentials were not corrected for liquid-liquid junction potentials.

### Analysis of intrinsic electrophysiological properties

Electrophysiological data were analysed using custom-made Python scripts, unless otherwise specified, using the pyABF (https://github.com/swharden/pyABF) and IPFX (https://github.com/AllenInstitute/ipfx) packages. Patch pipettes (4-6 MΩ) were pulled from standard-wall borosilicate capillaries and files with potassium-based internal solution (in mM): 140 K-gluconate, 10 KCl, 10 HEPES, 4 Na-Phosphocreatine, 4 ATP-Mg, 0.4 GTP-Na (pH 7.3, 290 mOsm). Biocytin (4g/mL) was finally added to the recording pipette.

Resting membrane potential (RMP) was measured immediately after whole-cell opening at injected current I = 0. Cells were considered spontaneously active when at least one action potential was observed in current-clamp mode without current injection.

For current-clamp recordings, holding current was adjusted to maintain membrane potential near −70 mV3. Input-output relationships were assessed using 250 ms current injections ranging from −200 to 1000 pA in 50 pA increments. Electrophysiological features were extracted using the pyABF package and IPFX package. Action potentials were detected when dV/dt exceeded 20 mV/s, with a minimum peak amplitude of −30 mV and a minimum threshold-to-peak amplitude of 2 mV. Spike threshold was defined at 5% of the average upstroke.

For each neuron, action potential firing curves were fitted with a sigmoid function: 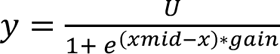, with U being the upper asymptote, xmid the current intensity evoking 50% of the upper asymptote and gain being the slope of the sigmoid. The fitting was done from the response to the lowest current intensity injected, to the last value evoking the highest firing frequency, to avoid any confounding effect due to saturation to the injected current. Spiking features were extracted from the rheobase sweep, which was determined injecting current from −30 pA until the first spike is evoked with 10 pA increments.

Passive membrane properties were assessed with 1s current injection from −250 to −100 pA, with 50 pA increments^36^. Sag amplitude was defined as (voltage peak – steady state voltage), with the voltage peak being the minimum value during the first 100ms of the current injection, and the steady state voltage being the mean voltage for the last 30ms of the current injection. Sag ratio was computed as 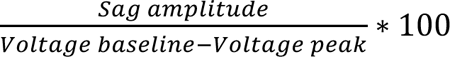, with the voltage baseline: the mean voltage during the last 100ms preceding current injection. Input resistance was defined as the coefficient of the linear fit between current-steps and the steady state voltages. The time constant tau was obtained from a single exponential fit of the voltage trace from current injection to the voltage peak using the following function: 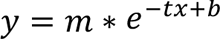, tau (in milliseconds) being defined as 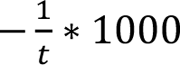.

Spontaneous excitatory postsynaptic currents (sEPSCs) were recorded in gap-free voltage-clamp mode at −70 mV over 20 s periods. Events were automatically detected and quantified using a custom MATLAB (MathWorks) script, then manually curated prior to analysis.

### Statistical analysis

All statistical analyses and graphical representations were performed using RStudio, MATLAB and GraphPad Prism (GraphPad Software, La Jolla, CA, USA). Statistical tests were selected according to sample size, data distribution, and variance homogeneity. Normality and homoscedasticity were evaluated prior to parametric analyses.

For RT–qPCR experiments, differences in gene expression between groups were assessed using unpaired two-tailed Student’s t-tests. Two-way ANOVA was used to evaluate the effects of genotype and cortical region or frequency band on EEG activity and electrophysiological intrinsic properties. When appropriate, post hoc multiple-comparison analyses were performed using corrected pairwise comparisons.

Parametric tests were used when normality and homoscedasticity assumptions were met; otherwise, when datasets did not satisfy assumptions for parametric, non-parametric or permutation-based approaches were applied.

Depending on cohort size, either exact permutation tests (small sample sizes) or Monte Carlo permutation simulations (9,999 iterations) were used to estimate null distributions and calculate significance values.

Kolmogorov–Smirnov (KS) tests were used to compare cumulative probability distributions of PNN mesh parameters and synaptic puncta measurements, including area and mean fluorescence intensity (Sup. Fig. 2 and 3). Repeated-measures correlation analyses were performed to assess relationships between synaptic marker intensity and local WFA intensity in synaptic puncta analyses.

In box-and-whisker plots, the central line represents the median, box boundaries indicate the interquartile range, and whiskers denote minimum and maximum values. Bar graphs are presented as mean ± standard error of the mean (SEM). Statistical significance was set at p < 0.05.

## RESULTS

### *Dlx5/6* inactivation in GABAergic neurons alters PNN-related gene expression

As transcription factors, *Dlx5* and *Dlx6* regulate gene expression through molecular mechanisms that remain poorly characterized in the adult brain. Given that *Dlx5/6* deletion profoundly affects PV interneuron density and cortical circuit function^8,11,12^, and the major role of PNNs as regulators of PV interneuron adult properties^13,37^, we investigated whether *Dlx5/6* inactivation in GABAergic neurons alters the expression of genes associated with PNN composition, assembly, and remodelling. We performed RT-qPCR analysis on tissue dissected from the parietal and prefrontal cortices of control and *Dlx5/6^VgatCre^* adult mice (P60) (Fig.1).

**Figure 1.**
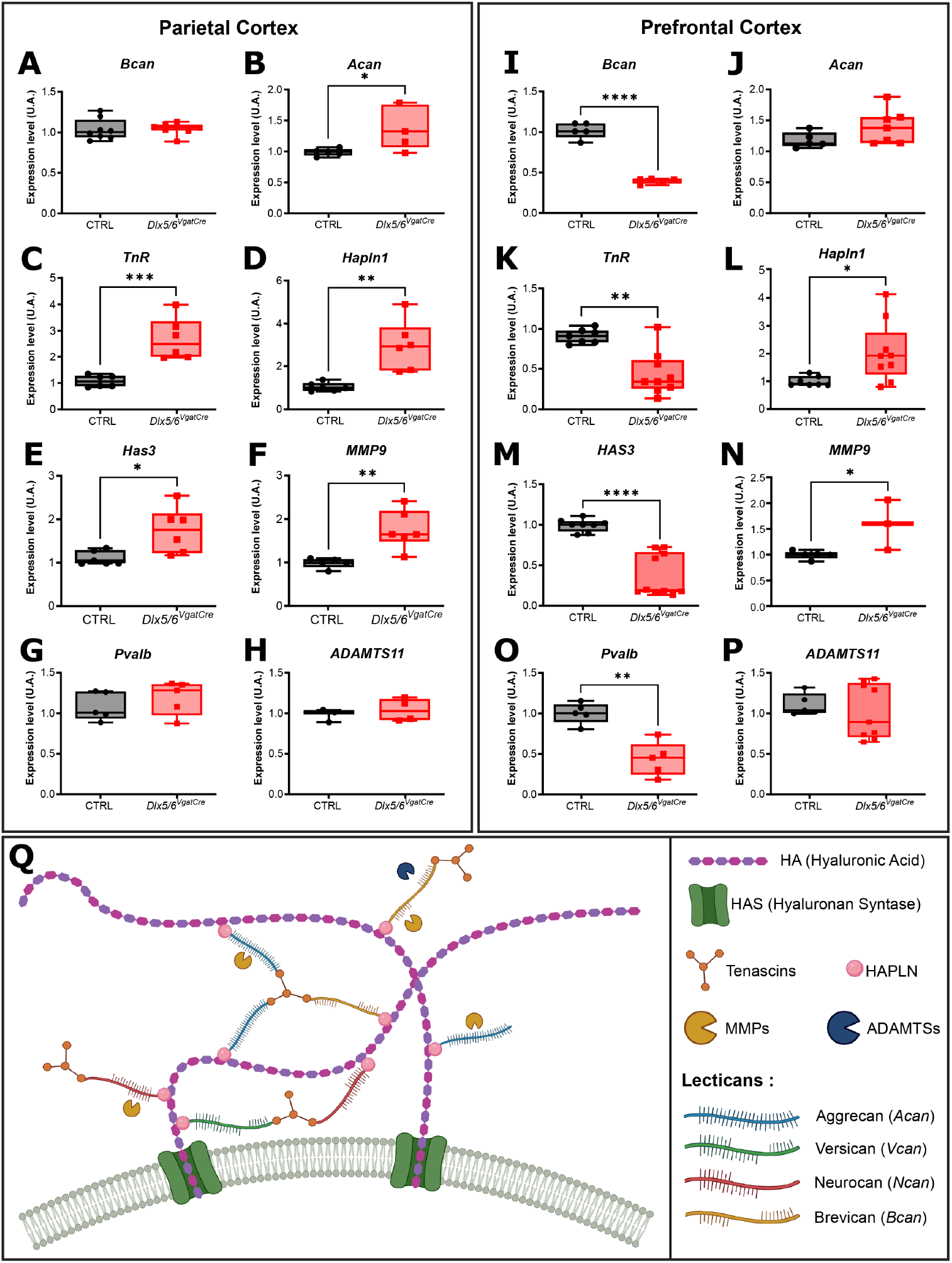
Region-specific transcriptional alterations of perineuronal net components following *Dlx5/6* inactivation in GABAergic neurons. Quantitative RT-PCR analysis of genes encoding perineuronal net (PNN) components and extracellular matrix-remodelling enzymes in the S1BF and PrL of adult control and *Dlx5/6^VgatCre^* mice. In the S1BF, *Dlx5/6* inactivation significantly upregulated expression of *Acan*, *Tnr*, *Hapln1*, *Has3* and *Mmp9* (**B-F**), whereas *Bcan*, *Pvalb* and *Adamts11* remained unchanged (**A, G, H**). In the PrL, a distinct transcriptional profile was observed, characterized by significant downregulation of *Bcan*, *Tnr*, *Has3* and *Pvalb* (**I, K, M, O**), alongside upregulation of *Hapln1* and *Mmp9* (**L, N**); *Acan* and *Adamts11* were not significantly affected (**J, P**). (**Q**) Schematic representation of PNN molecular architecture, illustrating the hyaluronic acid (HA) backbone synthesized by hyaluronan synthases (HAS), chondroitin sulphate proteoglycans (CSPGs; Aggrecan, Brevican, Versican and Neurocan), link proteins (HAPLN), Tenascins, and ECM-remodelling enzymes including matrix metalloproteinases (MMPs) and ADAMTS proteases. Data are presented as box-and-whisker plots (median, interquartile range; whiskers: min–max; *n* = 6 mice per group). Group comparisons were performed using two-tailed unpaired Student’s *t*-tests. \**p* < 0.05, \*\**p* < 0.01, \*\*\**p* < 0.001, \*\*\*\**p* < 0.0001.

In the parietal cortex*, Dlx5/6* inactivation induced a broad upregulation of several genes encoding structural and regulatory components of the PNN matrix (Fig. 1A–H). Expression of *Acan* (Aggrecan), a major chondroitin sulphate proteoglycan and core constituent of PNNs (Fig. 1Q)^38^, was significantly increased by 41% in *Dlx5/6^VgatCre^*mice (CTRL = 0.9878, *Dlx5/6^VgatCre^* = 1.395; *p* < 0.0348; Fig. 1B). Similarly, expression of *Tnr* (Tenascin-R) and *Hapln1* (Hyaluronan and Proteoglycan Link Protein 1), both implicated in PNN stabilization and matrix organization^26^, were respectively significantly elevated 2.75-fold (CTRL = 1.077, *Dlx5/6^VgatCre^* = 2.690; *p* = 0.0022; Fig. 1C) and 2.85-fold (CTRL = 1.040, *Dlx5/6^VgatCre^* = 2.965; *p* = 0.0022; Fig. 1D). Genes involved in extracellular matrix synthesis and remodelling were also upregulated, including *Has3* (Hyaluronan Synthase 3), which encodes the enzyme responsible for the synthesis of the hyaluronan backbone of PNNs^20^ and was increased by 57% (CTRL = 1.112, *Dlx5/6^VgatCre^* = 1.747; *p* = 0.0260; Fig. 1E), and *Mmp9* (Matrix Metalloproteinase 9) a protease involved in extracellular matrix degradation and remodelling^20^, which was increased by 80% (CTRL = 0.9632, *Dlx5/6^VgatCre^* = 1.732; *p* = 0.0043; Fig. 1F). Together, these findings indicate that *Dlx5/6* inactivation promotes the expression of multiple PNN-associated genes involved in both catabolism and anabolism of PNNs in the parietal cortex.

In contrast, the prefrontal cortex displayed a markedly different transcriptional profile (Fig. 1I–P). Expression of several structural PNN components was significantly reduced in *Dlx5/6^VgatCre^* mice, including *Bcan*, decreased by 60% (Brevican) (CTRL = 0.9914, *Dlx5/6^VgatCre^* = 1.5850; *p* = 0.0357; Fig. 1I) and *TnR,* decreased by 41% (CTRL = 0.9974, *Dlx5/6^VgatCre^* = 0.5904; *p* = 0.0006; Fig. 1K). However, *Hapln1* expression was significantly increased (2-fold; CTRL = 1.121, *Dlx5/6^VgatCre^* = 2.270; *p* = 0.0229; Fig. 1L). *Has3* expression was also significantly decreased, by 57% (CTRL= 0.9913, *Dlx5/6^VgatCre^*= 0.4276; *p* = 0.0034; Fig. 1M), whereas *Mmp9* levels were significantly increased (Fig. 1N), revealing a region-specific pattern of PNN-related transcriptional dysregulation.

To control for potential effects related to changes in PV interneuron density, we also quantified *Pvalb* expression. Consistent with our previous observations^8^, *Pvalb* expression was unchanged in the parietal cortex (Fig. 1G). In contrast, it was reduced by approximately 44% in the prefrontal cortex of the *Dlx5/6* recombined mice (CTRL= 1.001, *Dlx5/6^VgatCre^* = 0.4395; *p* = 0.0001; Fig.1O), consistent with the reduced density of PV^+^ neurons observed in this region. We additionally assessed the expression levels of other PNN-associated genes involved in PNN dynamic remodelling, which were similarly dysregulated in a region-specific manner (Sup. Fig. 1).

Together, these results demonstrate that *Dlx5/6* inactivation in GABAergic neurons induces pronounced, region-dependent transcriptional alterations in genes associated with PNNs structure and remodelling. These findings support a role for *Dlx5/6* in the molecular regulation of PV interneuron-associated extracellular matrix organization and suggest that altered PNN composition contributes to the functional deficits observed in *Dlx5/6^VgatCre^*cortical circuits.

### Dlx5/6 inactivation alters perineuronal net structure

We next examined whether the molecular changes induced by *Dlx5/6* inactivation in GABAergic neurons were accompanied by structural reorganization of PNNs. We first quantified the density of PV^+^ interneurons and WFA^+^ PNNs in the somatosensory barrel-field (S1BF) and prelimbic (PrL) cortices, two specific cortical territories of the parietal and prefrontal cortices, respectively, of adult (P60) control and *Dlx5/6^VgatCre^*mice (Sup. Figure 2). PNNs were visualised using Wisteria floribunda agglutinin (WFA)^+^ labelling, while PV^+^ interneurons were identified by immunofluorescence (Sup. Fig. 2A-B’).

Consistently with previous observations^8^, PV^+^ interneurons density was unchanged in the S1BF of *Dlx5/6^VgatCre^* mice relative to controls (Sup. Fig. 2C), whereas the PrL exhibited a significant 40% reduction of PV^+^ interneurons density in *Dlx5/6^VgatCre^* mice (Sup. Fig. 2D). Interestingly, WFA^+^ cell density was unaffected in both regions, suggesting that *Dlx5/6* inactivation does not alter overall PNN^+^ neurons density (Sup. Fig. 2D-C).

Given that RT-qPCR data revealed pronounced transcriptional alteration in genes involved in PNN composition and homeostasis, we investigated whether this invalidation disrupted PNN architecture rather than cell density.

We therefore performed a high-resolution structural analysis of WFA-labelled PNNs using the semi-automated image analysis pipeline APNU2.0 implemented in Fiji^33^. This approach enabled quantitative assessment of multiple parameters describing the size, shape, and fluorescence intensity of individual PNN mesh units in both 2D and 3D reconstructions. Analyses were conducted on immunofluorescent labelled cortical sections of adult control and *Dlx5/6^VgatCre^* mice (P60).

Quantitative analysis revealed significant structural remodelling of PNN mesh architecture in *Dlx5/6^VgatCre^* mice across both the PrL and S1BF regions (Figure 2).

**Figure 2.**
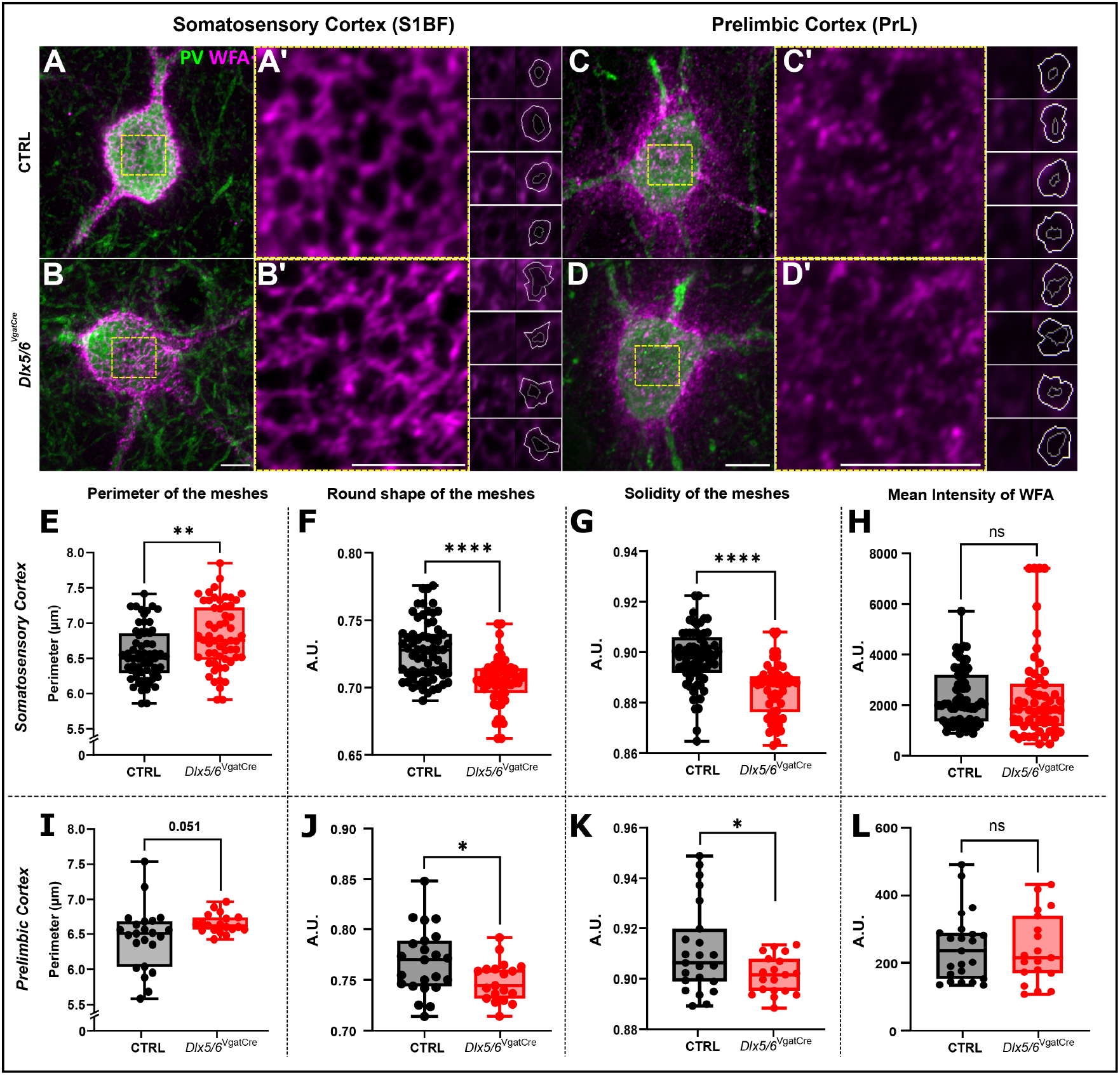
*Dlx5/6* inactivation induces region-specific remodelling of perineuronal net mesh architecture. Representative confocal images of WFA-labelled perineuronal nets (PNN^+^; magenta) surrounding Parvalbumin-positive (PV⁺; green) interneurons in the S1BF (**A-B’**) and PrL (**C-D’**) of control and *Dlx5/6^VgatCre^* mice. Enlarged views of the boxed regions are shown in **A’, B’, C’, D’**. Right panels illustrate representative mesh reconstructions generated using the APNU2.0 Fiji macro ^33^ for quantitative analysis of PNN architecture. In the S1BF, *Dlx5/6* inactivation significantly increased mesh perimeter (**E**) while reducing mesh roundness (**F**) and solidity (**G**), indicative of altered mesh geometry. Mean WFA fluorescence intensity was not significantly different between genotypes (**H**). In the PrL, mesh perimeter showed a trend toward increase (*p* = 0.051; I), whereas mesh roundness (**J**) and solidity (**K**) were significantly reduced in *Dlx5/6^VgatCre^* mice; mean WFA fluorescence intensity was unaffected (**L**). Mesh-based analyses included 2,763 meshes from 7 control mice and 3,160 meshes from 5 *Dlx5/6^VgatCre^* mice. Cell-based quantification was performed by averaging mesh parameters per individual WFA^+^ neuron; each data point represents one cell (S1BF: control, *n* = 51; *Dlx5/6^VgatCre^*, *n* = 44; PrL: control, *n* = 23; *Dlx5/6^VgatCre^*, *n* = 19). Data are presented as box-and-whisker plots (median, interquartile range; whiskers: min–max). Statistical significance was assessed using permutation tests (9,999 Monte Carlo permutations). \**p* < 0.05; \*\**p* < 0.01; \*\*\**p* < 0.001; \*\*\*\**p* < 0.0001; ns, not significant. Scale bars: 5 μm (**A-D’**); 2.5 μm (mesh reconstructions).

In the somatosensory cortex, *Dlx5/6* invalidated PV neurons formed a lattice-like PNN structure (Fig. 2A-B’), but displayed a significant increase in PNN mesh perimeter (Fig. 2E), indicating an enlarged mesh structure (CTRL = 6.55 µm; *Dlx5/6^VgatCre^* = 6.820; *p* = 0.031, Monte Carlo Permutation test, 9,999 permutations). *Dlx5/6^VgatCre^* mice also exhibited alterations in matrix organization, as reflected by changes in the solidity parameter, which describes PNN mesh compactness (Fig. 2G). Solidity values were significantly reduced in *Dlx5/6^VgatCre^*mice, from almost 3% (CTRL = 0.8978, *Dlx5/6^VgatCre^*= 0.8850; *p* < 0.0001), indicating a less compact extracellular matrix organization. In parallel, mesh roundness was decreased in the absence of *Dlx5/6* (Fig. 2F), revealing a shift towards a more irregular geometry (CTRL = 0.7272, *Dlx5/6^VgatCre^* = 0.7044; *p* < 0.0001). By contrast, overall, WFA fluorescence intensity remained comparable between genotypes (Fig. 2H).

In the prelimbic cortex, PNNs followed a slightly different morphological phenotype (Fig. 2C-D’). Invalidation of *Dlx5/6* induced alterations in mesh perimeter showed a strong trend toward an increase in *Dlx5/6^VgatCre^*mice from over 3.69% (CTRL = 6,4231, *Dlx5/6^VgatCre^* = 6,6598; *p* = 0.051, Monte Carlo Permutation test, 9,999 permutations), although this effect did not reach statistical significance. However, analysis of the cumulative distributions of individual mesh perimeter values using the Kolmogorov-Smirnov (KS) test revealed that control and mutant distributions were significantly different (*p* = 2.24×10^-05^), with a leftward shift in the mutant distribution indicating an overall increase in mesh perimeter (Sup. Fig. 2I).

Similarly to what was observed in the S1BF, solidity was significantly reduced in the absence of *Dlx5/6* (CTRL = 0.9084, *Dlx5/6^VgatCre^* = 0.8993; *p* < 0.0319; Fig. 2K), indicating decreased matrix compactness. Mesh roundness was also decreased in *Dlx5/6^VgatCre^*mice (Fig. 2J), consistent with altered PNN geometry. As observed in the somatosensory, WFA fluorescence intensity remained unchanged between groups.

To further characterize these alterations at the population level, we analysed the cumulative distributions of individual mesh parameters using Kolmogorov–Smirnov (KS) tests. For all analysed variables, cumulative distributions were significantly shifted in *Dlx5/6^VgatCre^*mice compared to CTRL (Sup. Fig. 2E-L), demonstrating widespread reorganization of PNN structural properties rather than isolated alterations.

Together, these results demonstrate that *Dlx5/6* inactivation induces profound remodelling of PNN architecture, affecting mesh size, geometry and matrix. These findings suggest that *Dlx5/6* primarily regulate PNN architecture without altering overall WFA-positive extracellular matrix density.

### Region-specific alterations in excitatory and inhibitory PNN-ensheathed soma synapses onto PV neurons

PNNs form a specialized extracellular scaffold surrounding the soma and proximal dendrites of PV interneurons, where they are intimately associated with excitatory and inhibitory perisomatic synapses. Through interactions with pre- and postsynaptic components, they contribute to the organization and stabilization of the perisomatic synaptic architecture^39^. Given the structural remodelling of PNNs observed in *Dlx5/6^VgatCre^* mice, we next examined whether inhibitory and excitatory perisomatic synapses surrounding PV interneurons were also remodelled in the somatosensory and prelimbic cortices.

Using high-resolution confocal microscopy, we quantified GABAergic (VGAT-positive) and Glutamatergic (VGLUT1-positive) presynaptic puncta associated with WFA-labelled PNNs surrounding PV interneuron somata in control and *Dlx5/6^VgatCre^* adult mice (P60) (Fig. 3A-D”).

**Figure 3.**
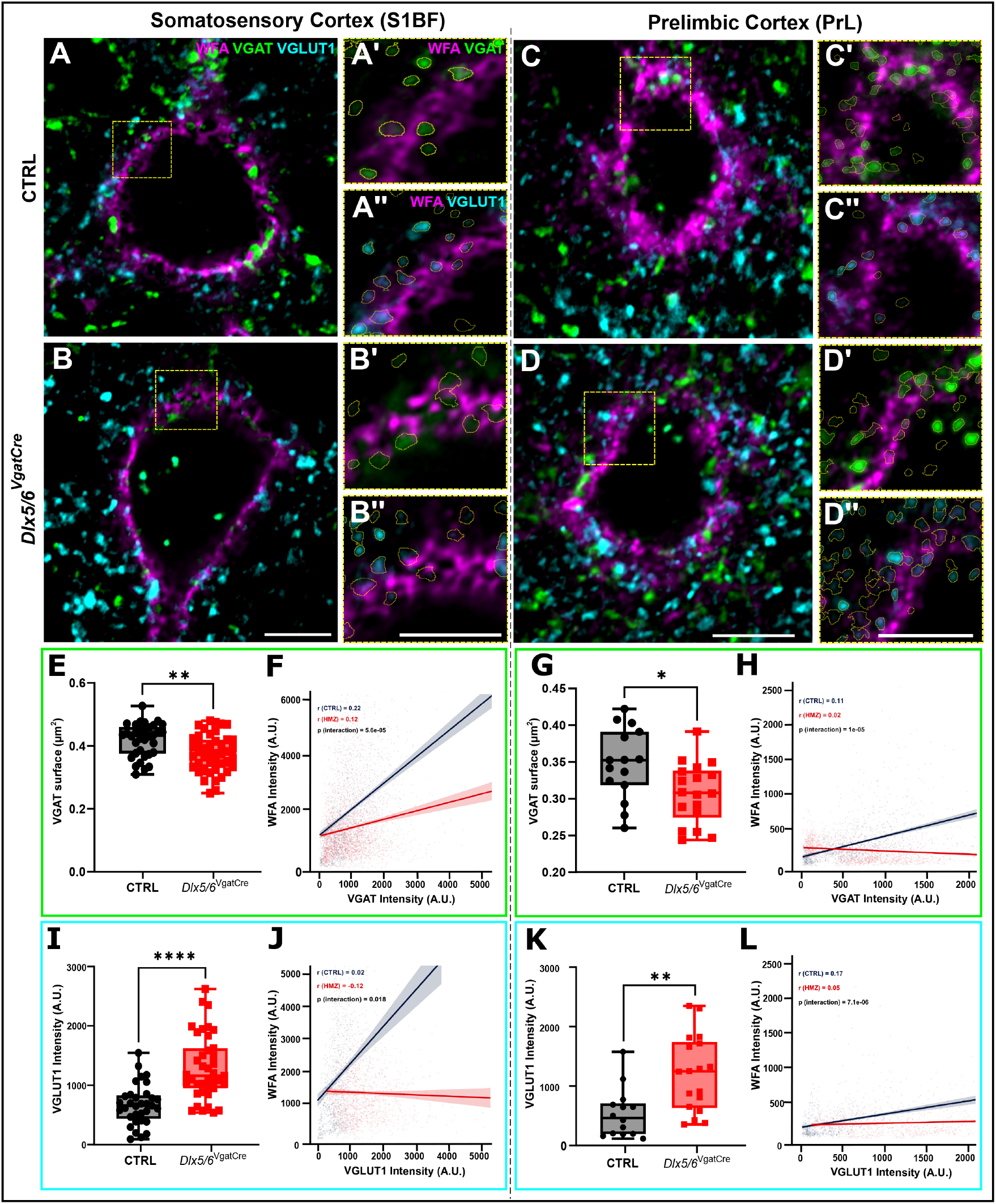
*Dlx5/6* inactivation alters the synaptic organization of inhibitory and excitatory inputs onto WFA-labelled perineuronal nets surrounding PV⁺ interneurons. Representative confocal images of Parvalbumin-positive (PV⁺) interneurons surrounded by WFA-labelled perineuronal nets (PNN^+^; magenta) in the S1BF (**A-A’’**) and PrL (**C-D’’**) of control and *Dlx5/6^VgatCre^* mice. Inhibitory presynaptic terminals were labelled with VGAT (green) and excitatory presynaptic terminals with VGLUT1 (cyan). Inserts (**A’-A’’, B’-B’’, C’-C’’, D’-D’’**) show higher-magnification views of the boxed regions, with WFA/VGAT and WFA/VGLUT1 channel combinations shown separately. Synaptic puncta were detected and quantified using circular reconstructions generated with the APNU2.0 Fiji macro^33^. In the S1BF, *Dlx5/6* inactivation significantly decreased VGAT puncta surface area (**E**), while the positive correlation between VGAT intensity and WFA intensity observed in controls was abolished in *Dlx5/6^VgatCre^* mice (**F**). VGLUT1 puncta intensity is significantlyincreased (**I**), and the positive correlation between VGLUT1 intensity and WFA intensity was similarly disrupted (**J**). In the PrL, VGAT puncta surface area was significantly reduced in *Dlx5/6^VgatCre^* mice (**G**), and the positive correlation between VGAT intensity and WFA intensity was lost (**H**). VGLUT1 puncta intensity was significantly increased (**K**), accompanied by a disruption of the positive WFA-VGLUT1 intensity correlation (**L**). For S1BF, VGAT analyses included 2,053 puncta from 34 cells across 5 control mice and 2,715 puncta from 42 cells across 8 *Dlx5/6^VgatCre^* mice; VGLUT1 analyses included 1,239 puncta from 34 cells across 5 controls and 2,296 puncta from 42 cells across 8 from *Dlx5/6^VgatCre^* mice. For PrL, VGAT analyses included 1,233 puncta from 15 cells across 5 control mice and 1,507 puncta from 18 cells across 8 *Dlx5/6^VgatCre^* mice; VGLUT1 analyses included 830 puncta from 15 cells across 5 controls and 794 puncta from 18 cells across 8 from *Dlx5/6^VgatCre^* mice. Correlation analyses (**F, H, J, L**) were performed using repeated-measures correlation with a group-by-predictor interaction term to assess genotype-dependent differences in regression slopes; ρ values are shown per genotype. Group comparisons were performed using Monte Carlo method (9,999 permutations). Quantifications were performed by averaging mesh parameters per individual WFA^+^ neuron; each data point represents one cell (S1BF: control, *n* = 34; *Dlx5/6^VgatCre^*, *n* = 42; PrL: control, *n* = 15; *Dlx5/6^VgatCre^*, *n* = 18). Data are presented as box-and-whisker plots (median, interquartile range; whiskers: min–max). \**p* < 0.05; \*\**p* < 0.01; \*\*\*\**p* < 0.0001. Scale bars: 10 μm (**A-D**); 5 μm (**A’-D’’**).

Analysis of VGAT-positive puncta revealed significant alterations in inhibitory synaptic organization in *Dlx5/6^VgatCre^* mice. In both cortical regions, VGAT puncta size was significantly reduced, by approximately 10% in the S1BF (Fig. 3E) and 12% in the PrL (Fig. 3G), indicating structural remodelling of inhibitory presynaptic contacts. VGAT puncta mean intensity did not differ significantly between genotypes in either region (Sup. Fig. 3I, K). However, cumulative distribution analysis of individual VGAT fluorescence values revealed significantly different distributions between genotypes in both regions (Sup. Fig. 3B, D), most pronounced in the PrL cortex (K.S. Test, *p* = 2.15 14e13 Sup. Fig. 3D), with a rightward shift of the mutant distribution indicating a tendency toward increased VGAT puncta fluorescence intensity in *Dlx5/6^VgatCre^* mice.

Excitatory presynaptic markers displayed the opposite profile. Indeed, VGLUT1-positive puncta showed significantly increased fluorescence intensity in both cortical regions (Fig.3 I, K), with increased by approximately 93% in the S1BF (CTRL = 673, *Dlx5/6^VgatCre^*= 1299; *p* < 0.0001), and an increase of 122% of intensity in the PrL (CTRL = 561.6, *Dlx5/6^VgatCre^* = 1247; *p* < 0.0001). This result could reflect changes in the number of presynaptic vesicles^40^.

In contrast, when puncta surface area was averaged per cell, no significant differences were observed between genotypes in either region (*p_S1BF_* = 0.6583; *p_PrL_* = 0.6959; Sup. Fig. 3M, O). However, cumulative distribution analysis revealed a slight rightward shift in VGLUT1 puncta surface area in the S1BF (Kolmogorov-Smirnov test, *p* = 0.123), suggesting a tendency to a smaller punctum, whereas a leftward shift was observed in the PrL (Kolmogorov-Smirnov test, *p* = 7.87×10^-4^), indicating an increase in the VGLUT1 puncta size.

These quantifications indicate that mutant mice display disrupted organization of synaptic puncta across cortical regions, consistent with a widespread reorganization of the perisynaptic environment rather than isolated synaptic alterations.

To further explore the relationship between synaptic organization and local PNN structure, we examined correlations between presynaptic marker intensity (VGAT or VGLUT1) and WFA fluorescence intensity at individual synaptic contacts. In control animals, presynaptic signal intensity positively correlated with local WFA intensity in both cortical regions, indicating a close association between synaptic properties and PNN organization (Fig. 3 F, H, J, L). Strikingly, this correlation was lost or largely weakened in *Dlx5/6^VgatCre^*mice. In the S1BF, the positive correlation between VGAT intensity and WFA intensities was reduced (ρCTRL = 0.22, ρ*Dlx5/6^VgatCre^* = 0.12, *p* = 5.6×10^-5^; Fig. 3F), as was the correlation involving VGLUT1 puncta (ρCTRL = 0.02, ρ*Dlx5/6^VgatCre^*= −0.12, *p* = 0.018; Fig. 3J). Similar reductions were observed in the PrL for VGAT puncta (ρCTRL = 0.11, ρ*Dlx5/6^VgatCre^* = 0.02, *p* = 1×10^-5^; Fig. 3H) and VGLUT1 puncta (ρCTRL = 0.17, ρ*Dlx5/6^VgatCre^* = 0.05, *p* = 7.1×10^-6^; Fig. 3L). Together, these results indicate a decoupling between extracellular matrix organization and synaptic architecture following *Dlx5/6* inactivation.

Collectively, these findings demonstrate that *Dlx5/6* inactivation in GABAergic neurons induces substantial remodelling of synaptic organization onto PNN-ensheathed PV interneurons. The combined alterations in inhibitory and excitatory presynaptic markers, together with the disruption of their association with local PNN structure, support a role for *Dlx5/6*-independent regulation of the extracellular matrix and in the stabilization and functional organization of cortical inhibitory circuits.

### *Dlx5/6* inactivation in GABAergic neurons alters intrinsic properties of PV neurons in somatosensory and prelimbic cortices

The observed alterations in PNN architecture and in the organisation of inhibitory and excitatory synaptic input onto PV interneurons prompted us to investigate whether these synaptic defects were accompanied by changes in the electrophysiological properties and connectivity of PV interneurons.

To address this question, we performed whole-cell patch-clamp recordings from layer V PV neurons in PrL and S1BF cortices of adult *Dlx5/6^VgatCre^* mice and control littermates (P67-P75). PV interneurons were visually identified for electrophysiological recordings using *AAV-PHP.eB/2-S5E2-HBB-chl-dTomato_2A_NLSdTomato* viral strategy that selectively labels Nav1.1-expressing fast-spiking interneurons^35^ (Fig. 4A), and were selected based on their characteristic fast-spiking electrophysiological profile^41^.

**Figure 4.**
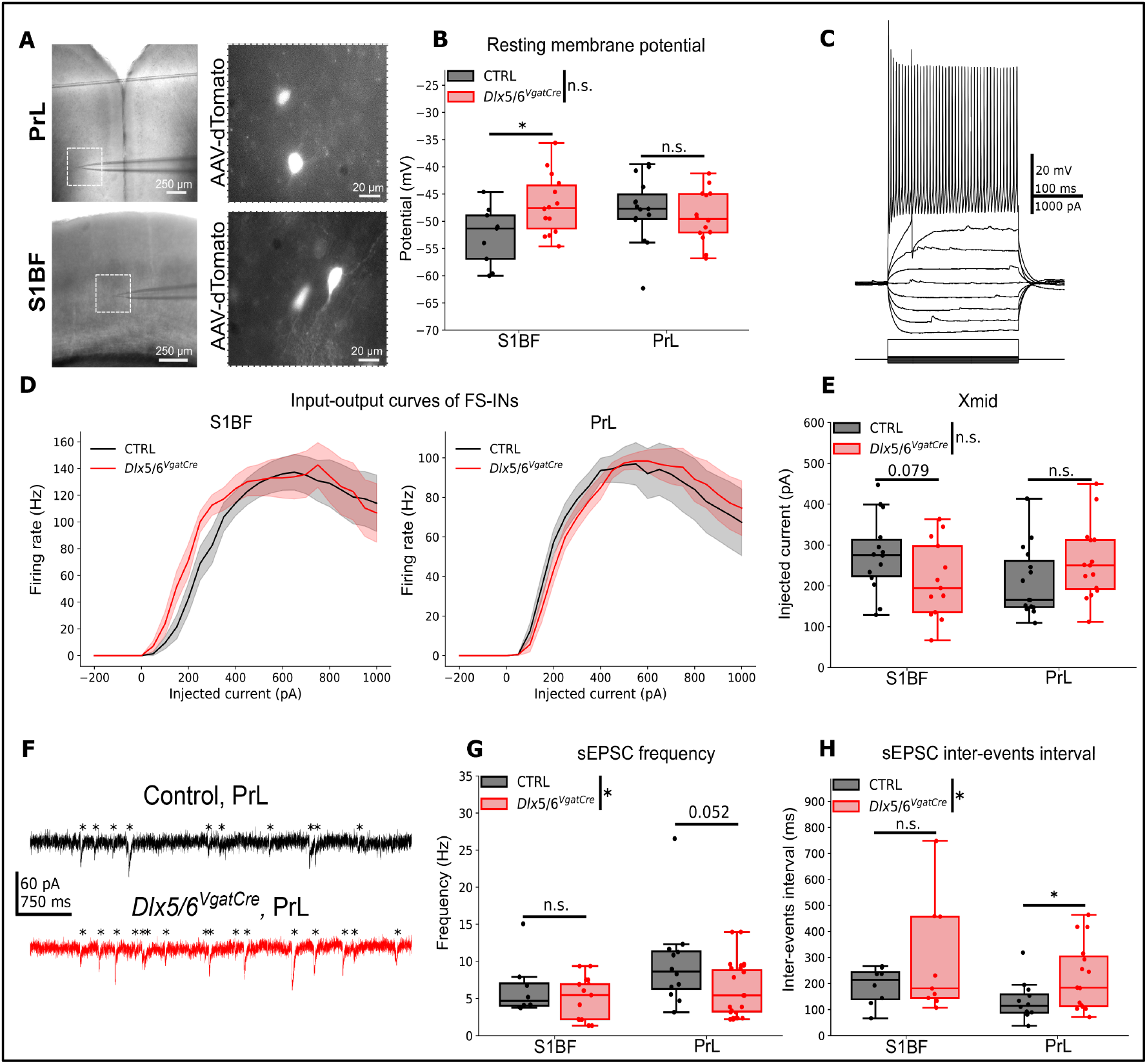
*Dlx5/6* inactivation in VGAT^+^ neurons alters intrinsic properties and spontaneous excitatory connectivity of fast-spiking interneurons. (**A**) Left: Representative images showing electrode positioning in prelimbic (PrL; top) and somatosensory (S1BF; bottom-up) cortices. Right: Representative fluorescence images from PrL (top) and S1BF (bottom) cortices. (**B**) Quantification of resting membrane potential. Two-way ANOVA revealed a significant interaction between the genotype and cortical region (*p* = 0.024917). Post-hoc within-region comparisons showed that *Dlx5/6*^VgatCre^ fast-spiking interneurons display a significantly higher resting membrane potential (*p* = 0.022679) in S1BF, but not in PrL. (S1BF: n_CTRL_ = 9, N_CTRL_ = 5; n*_Dlx5/6VgatCre_* = 14, N*_Dlx5/6VgatCre_* = 5; PrL: n_CTRL_ = 15, N_CTRL_ = 5; n*_Dlx5/6VgatCre_* = 14, N*_Dlx5/6VgatCre_* = 6). (**C**) Representative recordings illustrating an input-output curve. (**D**) Population input-output curves in S1BF (left) and PrL (right). Repeated-measure ANOVA revealed no significant effect of genotype, or genotype x current interaction in either cortical region. Dark lines represent population mean and shaded area the SEM. (**E**) Quantification of Xmid values derived from sigmoid fits of input-output curves. Two-way ANOVA revealed a significant genotype × cortex interaction, whereas no main effect of genotype or cortical region was detected. Post hoc within-region comparisons did not reveal significant differences between genotypes. (S1BF: p *=* 0.079; n_CTRL_ = 14, N_CTRL_ = 6; n*_Dlx5/6VgatCre_* = 13, N*_Dlx5/6VgatCre_* = 6). (**F**) Representative traces of spontaneous excitatory postsynaptic currents (sEPSCs) recorded from fast-spiking interneurons in the medial prefrontal cortex (PrL) in control mice (top) and *Dlx5/6*^VgatCre^ (bottom). * Indicates where the sEPSCs are in each representative recording. (**G**) Quantification of sEPSCs frequency. Two-way ranked ANOVA revealed a significant genotype effect. Within cortex comparison did not reveal differences between genotypes (PrL: n_CTRL_ = 12, N_CTRL_ = 5; n*_Dlx5/6VgatCre_* = 15, N*_Dlx5/6VgatCre_* = 6). (**H**) Quantification of sEPSC inter-events intervals. Two-way ranked ANOVA revealed a significant genotype effect. Within cortex comparison (S1BF: Permutation Test; PrL: t-test) showed significantly increased inter-events intervals in PrL fast-spiking interneurons from *Dlx5/6*^VgatCre^ mice (PrL: n_CTRL_ = 12, N_CTRL_ = 5; n*_Dlx5/6VgatCre_* = 15, N*_Dlx5/6VgatCre_* = 6). Statistical significance was assessed using unpaired two-tailed Student’s t-tests, Monte Carlo method (9,999 permutations) and Two-way ranked ANOVA. \**p* < 0.05.; n.s., not significant.

Analysis of fast-spiking intrinsic membrane properties revealed region-specific alterations in *Dlx5/6^VgatCre^* mice (Fig. 4). In the somatosensory cortex, fast-spiking interneurons from *Dlx5/6* recombined mice exhibited a significantly depolarized resting membrane potential (RMP) compared with controls (*p* = 0.023, CTRL: −52.69 mV; *Dlx5/6^VgatCre^*: −46.91 mV; Fig. 4B), while the spiking threshold remained unchanged (*p* = 0.741, CTRL: −48.03 mV; *Dlx5/6^VgatCre^*: −47.47 mV; Sup. Fig. 4C). This result suggests that mutant fast-spiking neurons rest closer to firing threshold than controls and may therefore be more readily excitable. However, the rheobase (*p =* 0.111, CTRL: 115.33 pA, *Dlx5/6^VgatCre^*: 80.77 pA; Sup. Fig. 4B) and input resistance were not altered (*p =* 0.204, CTRL: 97.79 MΩ; *Dlx5/6^VgatCre^*: 125.03 MΩ; Sup. Fig. 4D), suggesting that some compensation could occur to restore neuron’s excitability.

Nevertheless, the input-output curve revealed a trend towards a reduced action potential firing (x-mid, current producing 50% of the maximum firing frequency) in mutant neurons (Fig. 4E), although this effect did not reach a statistical significance (*p =* 0.079, CTRL: 276.34 pA; *Dlx5/6^VgatCre^*: 210.48 pA; Fig. 4G), suggesting altered excitability in S1BF fast-spiking interneurons.

In contrast, no significant alterations in intrinsic membrane properties were detected in fast-spiking interneurons from the prelimbic cortex (PrL) (Fig. 4B-E and Sup. Fig. 4B-F).

To evaluate the PV neuron local connectivity, we next examined the spontaneous excitatory postsynaptic currents (sEPSCs) in fast-spiking interneurons from both cortical regions. Analysis of sEPSC frequency using two-way ranked ANOVA revealed a significant genotype effect indicating an overall reduction in excitatory synaptic event frequency in *Dlx5/6^VgatCre^* mice (*p* = 0.0497, S1BF, CTRL: 6.34 Hz; *Dlx5/6^VgatCre^*: 5.01 Hz; PrL, CTRL: 9.73 Hz; *Dlx5/6^VgatCre^*: 6.02 Hz; Fig. 4G). Consistent with this observation, analysis of inter-event intervals revealed a significant genotype effect (*p* = 0.048, S1BF, CTRL: 191.25 ms; *Dlx5/6^VgatCre^*: 291.08 ms; PrL, CTRL: 132.63 ms; *Dlx5/6^VgatCre^*: 226.92 ms; Fig. 4H), reflecting increased intervals between excitatory synaptic events. Parametric t-test further demonstrated that PV interneurons from the PrL cortex of *Dlx5/6^VgatCre^* mice exhibited significantly increased inter-event intervals compared with controls (*p* = 0.026, Fig. 4H), indicative of reduced spontaneous excitatory drive, and a trend for reduced sEPSC frequency (*p* = 0.052; Fig. 4G). No difference in amplitude was observed.

Together, these findings demonstrate that *Dlx5/6* inactivation in GABAergic neurons induces region-specific electrophysiological alterations in PV interneurons, characterized by an increased resting membrane potential in the somatosensory cortex and reduced excitatory synaptic input in the prelimbic cortex. These alterations likely contribute to the dysfunction of inhibitory circuits, leading to impaired cortical activity in *Dlx5/6^VgatCre^* mice.

### *Dlx5/6* inactivation disrupts frontal cortex task-evoked gamma oscillations

Given the alterations in PV^+^ intrinsic properties and excitatory synaptic inputs observed in *Dlx5/6^VgatCre^* mice, we next investigated whether cortical network activity was affected.

Because gamma oscillations critically depend on the activity and synchronization of Parvalbumin interneurons^15,16^, we recorded EEG activity in adult *Dlx5/6^VgatCre^* mice and control littermates (P60–80) during a social interaction task, using fronto-parietal and parieto-cerebellar derivations, hereafter referred to as frontal and parietal recordings, respectively. Indeed, social interaction is known to engage PV-dependent gamma oscillations in the prefrontal cortex^12,42–44^.

Power spectral density was assessed during the active social interaction period and compared with baseline epochs recording during the same session, defined as periods of no active social interaction, for both derivations.

Analysis of baseline epochs revealed significant alterations in cortical oscillatory dynamics in *Dlx5/6^VgatCre^* mice. In frontal recordings, mutant mice exhibited reduced relative power in the Delta band (0.5-5 Hz; *p* = 0.0159; Permutation Test, n = 5 CTRL, 5 *Dlx5/6^VgatCre^*) together with increased Theta-band power (5-10 Hz; *p* = 0.0079; Permutation Test, n = 5 CTRL, 5 *Dlx5/6^VgatCre^*; Fig. 5C). Similar alterations were observed in parietal recordings, with a trend toward reduced Delta power (*p* =0.0548) and a marked increase in Theta-band power (*p* =0.00397; Permutation Test, n = 5 CTRL, 5 *Dlx5/6^VgatCre^*; Fig.5A), indicating *Dlx5/6* deficiency constitutively alters cortical network oscillatory activity.

**Figure 5.**
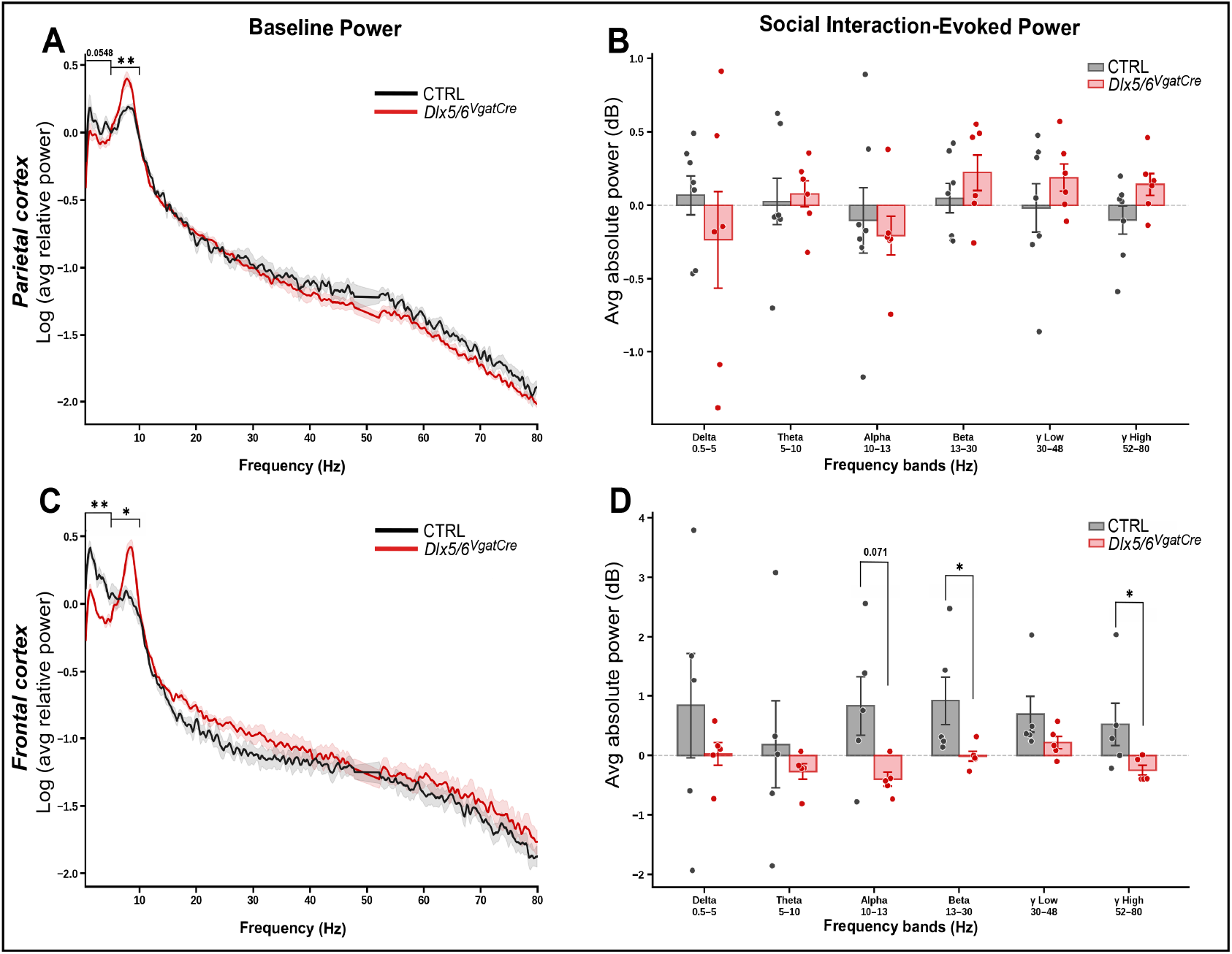
*Dlx5/6^VgatCre^*mice display altered baseline oscillatory activity and impaired task-evoked prefrontal gamma oscillations. (**A, C**) Log-transformed averaged relative power spectra recorded from the parietal cortex (**A**) and frontal cortex (**C**) of control (black) and *Dlx5/6^VgatCre^* mice (red) during baseline recordings. Dark lines represent population mean and shaded areas represent mean ± SEM. (**A, C**). For each animal, power spectra were normalized to the total spectral power across the 0–80 Hz frequency range, excluding the 48–52 Hz band. Relative to controls, *Dlx5/6^VgatCre^*mice exhibited reduced delta-band oscillatory power (0.5-5 Hz) together with increased theta-band activity (5-10 Hz). (**B, D**) Quantification of task-evoked absolute power across frequency bands (Delta: 0.5-5 Hz; Theta: 5-10 Hz; Alpha: 10-13 Hz; Beta: 13-30 Hz; Low Gamma: 30-48 Hz; High Gamm: 52-80 Hz) during social interaction task. In the prefrontal cortex (**D**) *Dlx5/6^VgatCre^* mice exhibited significantly reduced Beta- and High Gamma-band power relative to controls, whereas lower-frequency bands were not significantly affected. In contrast, no significant differences in social interaction-evoked power were detected in the parietal cortex (**B**), indicating that Gamma-band deficits are specific to the prefrontal cortex and emerge selectively during social interaction-evoked network recruitment. Statistical significance was assessed using exact permutation tests (252 permutations). Control, *n* = 5 mice; *Dlx5/6^VgatCre^*, *n* = 5 mice. *p < 0.05; **p < 0.01.

To specifically assess task-evoked oscillatory responses, we quantified for each animal the change in power between social interaction relative to baseline periods at each frequency bin.

During social interaction, control mice displayed a broad increase in frontal cortical power across multiple frequency bands (Fig. 5D). In contrast, *Dlx5/6^VgatCre^* mice exhibited impaired task-evoked oscillatory responses, with a significant reduction in high Gamma frequency range (52-80 Hz) in the frontal cortex compared to controls (*p* = 0.04; Permutation test; CTRL=0.52; *Dlx5/6^VgatCre^* =-0.25), indicating deficient recruitment of fast oscillations during social interaction. A similar reduction was observed in the Beta frequency range (13-30 Hz; *p* = 0.032; CTRL=0.92; *Dlx5/6^VgatCre^* = −0.01; Fig. 5D).

In contrast, no differences in task-evoked oscillatory activity were detected in parietal recordings (Fig. 5B). These findings suggest that network recruitment deficits in *Dlx5/6^VgatCre^* are specific to the frontal cortex and to social interaction-related network recruitment.

Together, these results demonstrate that *Dlx5/6^VgatCre^*mice fail to appropriately recruit frontal Gamma oscillations during social interaction. Given that Gamma-band activity emerges from the coordinated activity of PV interneuron networks, these findings suggest that the reduction in PV interneuron density and the functional alterations of surviving PV neurons contribute to impaired dynamic recruitment of cortical circuits during social processing.

## DISCUSSION

The present study highlights a novel role for *Dlx5/6* in the maintenance of mature cortical inhibitory networks, an aspect that has remained poorly characterized due to the embryonic lethality associated with conventional knockout models^11,12^. Although *Dlx5/6* are well established as regulators of GABAergic interneuron development^9,10^, our findings demonstrate that their activity remains essential after cortical development has been completed. By integrating molecular, structural, electrophysiological and network-level analyses, we show that conditional *Dlx5/6* inactivation disrupts a coordinated program linking extracellular matrix architecture, perisomatic synaptic organization, and cortical synchronization. Importantly, these alterations occur despite only modest changes in PV interneuron density, indicating that the adult function of *Dlx5/6* is not primarily to preserve interneuron number, but to maintain the specialized microenvironment required for mature PV interneuron networks to function normally. This distinction is evident across all levels of analysis. While PV cell density was only modestly affected in the prelimbic cortex and largely preserved in the somatosensory cortex (Sup. Fig.2)^8^, pronounced alterations were observed in PNN architecture, perisomatic synaptic organization, intrinsic membrane properties and gamma oscillations both in the prelimbic and somatosensory cortices. Since PNNs are an essential partner of PV-neurons functional integrity^20–22,27^, our results support the notion that developmental transcription factors such as *Dlx5/6* continue to play essential roles in maintaining circuit function long after cortical maturation has been completed.

Our molecular analyses suggest that *Dlx5/6* contribute to the homeostasis of the extracellular matrix surrounding PV interneurons. Rather than affecting a single component, *Dlx5/6* inactivation disrupted the coordinated expression of multiple genes involved in PNNs assembly, stabilization and remodelling, including *Acan*, *Tnr*, *Hapln*1, *Has3* and *Mmp9*. Interestingly, the transcriptional responses differed substantially between the parietal and prefrontal cortices. Such divergence is unlikely to arise only by differences in neuronal activity^45,46^, as PNN composition and molecular organization are known to vary considerably across cortical regions^47,48,20^. Condensed pericellular matrix assembly requires four essential components: hyaluronan synthetized by Has3, a CSPG of the lectican family (primarily Aggrecan), a link protein (such as HAPLN1), and Tenascin-R^20,49,50^. Inactivation studies have investigated the role of individual PNN components in matrix homeostasis and their physiological consequences^20,26^. Among the lectican family, Aggrecan is essential for PNN formation; its absence prevents the assembly of PNNs^38^. Other lecticans, including Neurocan, Versican, and Brevican, are present in most PNNs, but their absence does not prevent PNN formation and aggregation^51^. Brevican is specifically enriched at the synaptic cleft, where it has been shown to facilitate excitatory synaptic transmission onto target cells^52^.

The consequences of overexpressing PNN-related genes have not been studied to the same extent. Evidence from the literature suggests that increased expression of structural components would promote the formation of a more condensed and stabilized matrix^53^.

Strikingly, following *Dlx5/6* inactivation, the parietal cortex and prefrontal cortex displayed opposing transcriptional responses (Fig. 1). In the parietal cortex, several genes involved in PNN assembly and stabilization were upregulated, including *Acan*, *Tnr*, *Hapln1*, and *Has3* (Fig. 1A-H). Although this pattern would be expected to yield a denser matrix^53^, structural analysis instead revealed enlarged, less circular and less compact PNN meshes (Fig. 2E–G), that do not correspond to increased matrix density. In contrast, the prefrontal cortex exhibited reduced expression of multiple structural components such as *Bcan, TnR and Has3* (Fig. 1I-P), associated with decreased PNN mesh circularity and compactness (Fig. 2J-K), which is consistent with a loss of matrix condensation.

Despite the opposite transcriptional response on PNN components and remodelling enzymes, the phenotypes in the two regions converge on a common structural phenotype: a loss of PNN mesh compactness and geometric regularity reflects a dysregulation of the homeostatic dynamic balance between PNN assembly and degradation^28,54^.

In the S1BF, the massive increase in *Mmp9* expression (+80%, Fig. 1F) suggests an “hyper-remodelling” state in which the concurrent upregulation of structural PNN components (*Acan, Tnr, Hapln1*; Fig. 1B-D) and their synthesis enzyme (*Has3*; Fig. 1E) is insufficient to offset this exacerbated enzymatic degradation, resulting in the same lack of condensation and structural instability observed in the PrL, where *Mmp9* upregulation (Fig. 1N) is associated with a decrease in PNN components (*Bcan, Tnr*; Fig. 1I, K) and their synthesis enzyme (*Has3*; Fig. 1M).

This result suggests that *Dlx5/6* do not act as simple activators of matrix synthesis, but rather as regulators of the homeostatic balance essential for PNN stability. This model of convergence underscores that PNN functional integrity in the adult brain depends not only on the availability of components but on the precision of the turnover dynamics orchestrated by the *Dlx5/6* transcriptional program.

These PNN architectural alterations converge toward similar structural and physiological phenotypes, suggesting a common biological function attained through distinct regional molecular mechanisms. *Dlx5/6* deficiency induced marked remodelling of PNN geometry in the adult cortex, affecting mesh perimeter, compactness and circularity. These morphological changes and the density analysis point toward an altered organization of the extracellular matrix rather than a degeneration of the structures themselves. This distinction is functionally relevant, given that PNNs are dynamic regulators of neuronal excitability, synaptic stability and cortical plasticity rather than static protective coats surrounding mature neurons^28^, and that these functions are highly sensitive to alterations in matrix composition or reorganization^27,36,53^.

One of the most conceptually significant observations emerging from this study is that PNN remodelling was accompanied by a disruption of the normal relationship between extracellular matrix and perisomatic presynaptic organization.

In control animals, we observed a robust positive correlation between local PNN density and the intensities of both excitatory (VGLUT1^+^; Fig. 3H, L) and inhibitory (VGAT^+^; Fig. 3F, J) presynaptic markers, indicating that extracellular matrix organization and synaptic organization are tightly coordinated at the perisomatic level. It suggests that the matrix acts as a precise scaffold that stabilizes synaptic inputs. Strikingly, *Dlx5/6* inactivation causes a decoupling of this relationship (Fig. 3H, L, F, J). The altered relationship between local WFA organization and presynaptic markers suggests that Dlx5/6 loss disrupts the coordinated organization of the PNN–synapse microenvironment. This disruption may contribute to impaired recruitment of PV interneuron networks. We propose that the loss of matrix condensation, driven by region-specific transcriptional dysregulation, leads to a destabilization of the synapse, ultimately explaining the failure of PV interneurons to synchronize cortical networks and support complex social behaviors.

At the level of individual presynaptic markers, *Dlx5/6* inactivation induced distinct alterations, similar in both cortical regions. VGLUT1 puncta displayed increased fluorescence intensity (Fig. 3I, K), which has previously been associated with more synaptic vesicles within excitatory terminals^55,56^, are indicative of an altered excitatory drive onto PV interneurons. This interpretation is supported by patch-clamp recordings from the prefrontal cortex, which showed reduced spontaneous excitatory activity of PV neurons (Fig. 4F-H). Alongside these observations, VGAT puncta exhibited reduced size in both cortices (Fig.3 E, G), suggesting a reorganization of inhibitory presynaptic elements. These findings, paired with excitatory alterations, support a broader dysregulation of the perisomatic synaptic balance surrounding PV interneurons^57^.

The electrophysiological phenotype of *Dlx5/6^VgatCre^*mice supports an uncoupling between synaptic strength and perineuronal stability, and cortex-specific adaptation. PV interneurons displayed quite different region-specific phenotypes. In the somatosensory cortex, we observed an intrinsic more depolarized resting membrane potential and a trend toward reduced Xmid (Fig. 4B, E), which suggests increased excitability although the rheobase was not affected (Sup. Fig. 4B). Comparable alterations of fast-spiking interneuron properties have been reported following chABC enzymatic PNN degradation in the barrel field cortex^36^, supporting a structural remodelling of the perineuronal matrix that directly impacts PV interneuron excitability.

By contrast, in the prelimbic cortex, the reduction in spontaneous excitatory drive onto PV interneurons likely compromises their rapid recruitment during behaviourally-relevant activity (Fig. 4F-H). Consistent with this reduced excitatory drive, the alterations observed in high Gamma power (52-80Hz, Fig. 5B) during the social interaction task in *Dlx5/6^VgatCre^* mice may similarly reflect a reduced recruitment of local inhibitory circuits to generate the high-frequency network oscillations required for normal behaviour. The PV-neurons-dependent frontal cortex Gamma oscillations (30-80 Hz) are essential for high-order cognitive functions, including sensory perception^58,59^, behavioural processes^60,61^ and social interactions^43,44^. These rhythms critically depend on the synchronous, high-frequency firing of PV interneurons^12^. Prior work showed that constitutive *Dlx5/6* heterozygous inactivation leads to gamma-band defects in the frontal cortex associated with social/cognitive phenotypes^12^. Together, our findings indicate that relatively subtle alterations in the organization and synaptic integration of PV interneurons are sufficient to impair the dynamic recruitment of cortical synchrony during behaviour, highlighting the importance of *Dlx5/6* for preserving the functional integrity of adult inhibitory networks.

We also observed altered Delta and Theta baseline power in both frontal and parietal EEG recordings (Fig. 5A, C), independently of behavioural tasks, suggesting that these deficits are intrinsic to *Dlx5/6* inactivation. Delta and Theta rhythms derive from network oscillations generated by interactions between cerebral structures, Theta rhythms being particularly dependent on septo-hippocampal circuitry, whereas Delta bands reflect slower cortico-thalamic network synchronization^62^. As *Dlx5/6* inactivation affects all GABAergic neurons throughout the telencephalon, the observed alterations in Delta and Theta oscillations likely arise from disturbances in GABAergic neurons activity implicated in these regions.

*Parvalbumin* expression and perineuronal net density are dynamic features of the adult cerebral cortex, exhibiting measurable fluctuations over the course of a single day^28^. Both markers remain sensitive to neuronal activity, as chronic reductions in cortical activity induced by sensory deprivation or pharmacological manipulations decrease *PV* expression and promote PNN remodelling^28,30^, whereas sustained network activity stabilizes PNNs and preserves the molecular and electrophysiological properties of PV interneurons^52,63^. Similarly, *Dlx5/6* expression is maintained under dynamic regulation in the adult brain. We recently showed that *Dlx5/6* expression is controlled by the activity-dependent neurotrophin BDNF^64^, consistent with the concept that neuronal activity continuously shapes the transcriptional programs underlying neuronal identity and cortical function^65^. Together, these findings suggest that coordinated changes in *Dlx5/6* expression, PV levels, and PNN density constitute physiological mechanisms through which cortical circuits are influenced by cerebral activity. Alterations in PV interneurons, Gamma oscillations and PNN integrity have each been involved in neuropsychiatric disorders such as schizophrenia, bipolar disorders^66^, autism spectrum disorders, among other conditions affecting cognition and social behaviour^29,67,68^. In post-mortem brains of patients with schizophrenia, reduced interneuron density has been reported in the prefrontal and cingulate cortices^69^, alongside a decreased PNN density in the amygdala and entorhinal cortex^70,71^; similarly, reduced PV interneuron density has been identified in the prefrontal cortex in autism^72^. Patients with schizophrenia also display altered prefrontal Gamma oscillations^73^ during working-memory tasks^74,75^. Rather than representing independent pathological features, our findings suggest that part of these abnormalities may arise from the disruption of a common organizational framework dedicated to the maintenance of PNN integrity, which underlies mature inhibitory circuits. Furthermore, *Dlx5/6* may constitute key transcriptional factors involved in the pathophysiology of neuropsychiatric disorders associated with GABAergic dysfunction^17^.

Several limitations should be acknowledged. Although the present study establishes strong associations between extracellular matrix remodelling, altered synaptic organization, PV interneuron physiology and cortical oscillations, it does not directly demonstrate causal relationships between these phenomena. Likewise, the transcriptional analyses cannot distinguish between direct *Dlx5/6* targets and secondary downstream responses. Future studies combining genomic approaches, such as cell-type-specific or single-nuclei approaches, targeted manipulation of extracellular matrix components, and inducible *Dlx5/6* deletion restricted to adulthood, would be required to define these mechanisms more precisely and confirm their adult-specific functional role, even though PNNs forms postnatally after cortical maturation.

Our findings identify a Dlx5/6-dependent program required for the adult organization and functional integrity of PV interneuron–PNN microcircuits. By coordinating extracellular matrix homeostasis, peri-somatic synaptic architecture and intrinsic neuronal properties, this program maintains the conditions for efficient cortical synchronization during behaviour. More broadly, our study supports the idea that developmental transcription factors continue to shape adult brain function, not only by specifying neuronal identity, but by maintaining the structural and physiological organization of mature neuronal microcircuits.

## ACKNOWLEDGMENTS

This research was supported in part by the ANR grant METABRAIN (ANR-21-CE14-0072) awarded to G.L., the Fondation NRJ–Institut de France grant (N° 216612), the Sorbonne Université Emergence grant “DEDUCE,” and the MNHN ATM grants “Relation” and “DlxUp” awarded to N.N.N. L.B. was supported by a doctoral grant from the Muséum national d’Histoire naturelle. We thank Xavier Marques and Cyril Willig for their technical support with confocal microscopy at the Centre de Microscopie et d’Imagerie Numérique (CeMIM), and Patricia Wils for her assistance with image analysis. We are grateful to Dr. Alexander Dityatev and Dr. Rahul Kaushik for generously sharing the modified APNU2.0 ImageJ macro, and deeply thankful to Dr. Valentin Ritou for his assistance with EEG recording analysis. We also thank the animal facility team for mouse care, and particularly Stéphane Sosinsky and Fabien Uridat. We are grateful to Aïcha Bennana and Lanto Courcelaud for their administrative assistance.

## Author Contributions

Conceptualization: G.L., N.N.-N.; Methodology: L.B., F.H., V.F., M.B.P., N.N.-N.; Validation: L.B., R.A., B.A., N.N.-N., G.L.; Formal analysis: L.B., R.A., B.A.; Investigation: L.B., R.A., B.A., F.H., P.D., L.C., A.F.; Resources: A.F., V.F., M.B.P.; Data curation: L.B., R.A., B.A.; Writing-original draft preparation: L.B., G.L., N.N.-N.; Writing-review and editing: all authors; Visualization: L.B., B.A.; Supervision: N.N.-N.; Project administration: N.N.-N.; Funding acquisition: G.L. and N.N.-N. All authors have read and agreed to the published version of the manuscript.

## CONFLICT OF INTEREST

The authors declare no conflict of interest.

## Notes

### Competing Interest Statement

The authors have declared no competing interest.

## REFERENCES

1. Isaacson, J. S. & Scanziani, M. How Inhibition Shapes Cortical Activity. Neuron 72, 231–243 (2011).

2. Houghton, G. & Tipper, S. P. Inhibitory mechanisms of neural and cognitive control: applications to selective attention and sequential action. Brain Cogn 30, 20–43 (1996).

3. Xu, S. et al. Neural Circuits for Social Interactions: From Microcircuits to Input-Output Circuits. Front Neural Circuits 15, 768294 (2021).

4. Garcia-Junco-Clemente, P., Tring, E., Ringach, D. L. & Trachtenberg, J. T. State-Dependent Subnetworks of Parvalbumin-Expressing Interneurons in Neocortex. Cell Rep 26, 2282–2288.e3 (2019).

5. Ferguson, B. R. & Gao, W.-J. PV Interneurons: Critical Regulators of E/I Balance for Prefrontal Cortex-Dependent Behavior and Psychiatric Disorders. Front Neural Circuits 12, 37 (2018).

6. de Lombares, C. et al. Dlx5 and Dlx6 expression in GABAergic neurons controls behavior, metabolism, healthy aging and lifespan. Aging (Albany NY) 11, 6638–6656 (2019).

7. Levi, G. et al. DLX5/6 GABAergic Expression Affects Social Vocalization: Implications for Human Evolution. Mol Biol Evol 38, 4748–4764 (2021).

8. Aouci, R. et al. Dlx5/6 Expression Levels in Mouse GABAergic Neurons Regulate Adult Parvalbumin Neuronal Density and Anxiety/Compulsive Behaviours. Cells 11, 1739 (2022).

9. Panganiban, G. & Rubenstein, J. L. R. Developmental functions of the Distal-less/Dlx homeobox genes. Development 129, 4371–4386 (2002).

10. Rubenstein, J. L., Nord, A. S. & Ekker, M. DLX genes and proteins in mammalian forebrain development. Development 151, dev202684 (2024).

11. Wang, Y. et al. Dlx5 and Dlx6 regulate the development of parvalbumin-expressing cortical interneurons. J Neurosci 30, 5334–5345 (2010).

12. Cho, K. K. A. et al. Gamma Rhythms Link Prefrontal Interneuron Dysfunction with Cognitive Inflexibility in Dlx5/6+/− Mice. Neuron 85, 1332–1343 (2015).

13. Druga, R., Salaj, M. & Al-Redouan, A. Parvalbumin - Positive Neurons in the Neocortex: A Review. Physiol Res 72, S173–S191 (2023).

14. Buzsáki, G. & Wang, X.-J. Mechanisms of Gamma Oscillations. Annu Rev Neurosci 35, 203–225 (2012).

15. Sohal, V. S., Zhang, F., Yizhar, O. & Deisseroth, K. Parvalbumin neurons and gamma rhythms enhance cortical circuit performance. Nature 459, 698–702 (2009).

16. Cardin, J. A. et al. Driving fast-spiking cells induces gamma rhythm and controls sensory responses. Nature 459, 663–667 (2009).

17. Marín, O. Interneuron dysfunction in psychiatric disorders. Nat Rev Neurosci 13, 107–120 (2012).

18. Filice, F., Janickova, L., Henzi, T., Bilella, A. & Schwaller, B. The Parvalbumin Hypothesis of Autism Spectrum Disorder. Front. Cell. Neurosci. 14, (2020).

19. Hu, H., Gan, J. & Jonas, P. Interneurons. Fast-spiking, parvalbumin+ GABAergic interneurons: from cellular design to microcircuit function. Science 345, 1255263 (2014).

20. Fawcett, J. W., Oohashi, T. & Pizzorusso, T. The roles of perineuronal nets and the perinodal extracellular matrix in neuronal function. Nat Rev Neurosci 20, 451–465 (2019).

21. Wingert, J. C. & Sorg, B. A. Impact of Perineuronal Nets on Electrophysiology of Parvalbumin Interneurons, Principal Neurons, and Brain Oscillations: A Review. Front. Synaptic Neurosci. 13, (2021).

22. Burket, J. A., Webb, J. D. & Deutsch, S. I. Perineuronal Nets and Metal Cation Concentrations in the Microenvironments of Fast-Spiking, Parvalbumin-Expressing GABAergic Interneurons: Relevance to Neurodevelopment and Neurodevelopmental Disorders. Biomolecules 11, 1235 (2021).

23. Testa, D., Prochiantz, A. & Di Nardo, A. A. Perineuronal nets in brain physiology and disease. Seminars in Cell & Developmental Biology 89, 125–135 (2019).

24. Tewari, B. P. et al. Perineuronal nets support astrocytic ion and glutamate homeostasis at tripartite synapses. Res Sq rs.3.rs-2501039 (2023) doi:10.21203/rs.3.rs-2501039/v1.

25. Dityatev, A. & Rusakov, D. A. Molecular signals of plasticity at the tetrapartite synapse. Curr Opin Neurobiol 21, 353–359 (2011).

26. Jakovljević, A. et al. Structural and Functional Modulation of Perineuronal Nets: In Search of Important Players with Highlight on Tenascins. Cells 10, 1345 (2021).

27. Testa, D., Prochiantz, A. & Di Nardo, A. A. Perineuronal nets in brain physiology and disease. Seminars in Cell & Developmental Biology 89, 125–135 (2019).

28. Harkness, J. H. et al. Diurnal changes in perineuronal nets and parvalbumin neurons in the rat medial prefrontal cortex. Brain Struct Funct 226, 1135–1153 (2021).

29. Huang, Y.-F. et al. The role of the perineuronal net in pathological mechanisms of neurological and psychiatric disorders. Brain awag058 (2026) doi:10.1093/brain/awag058.

30. Devienne, G. et al. Regulation of Perineuronal Nets in the Adult Cortex by the Activity of the Cortical Network. J. Neurosci. 41, 5779–5790 (2021).

31. Miyata, S. & Kitagawa, H. Chondroitin 6-Sulfation Regulates Perineuronal Net Formation by Controlling the Stability of Aggrecan. Neural Plasticity 2016, 1305801 (2016).

32. The Mouse Brain in Stereotaxic Coordinates, Compact - Edition 3 - By George Paxinos and Keith B.J. Franklin Elsevier Educate. https://www.educate.elsevier.com/book/details/9780123742445.

33. Kaushik, R. et al. Fine structure analysis of perineuronal nets in the ketamine model of schizophrenia. European Journal of Neuroscience 53, 3988–4004 (2021).

34. Strackeljan, L. et al. Microglia Depletion-Induced Remodeling of Extracellular Matrix and Excitatory Synapses in the Hippocampus of Adult Mice. Cells 10, 1862 (2021).

35. Vormstein-Schneider, D. et al. Viral manipulation of functionally distinct interneurons in mice, non-human primates and humans. Nat Neurosci 23, 1629–1636 (2020).

36. Chu, P. et al. The Impact of Perineuronal Net Digestion Using Chondroitinase ABC on the Intrinsic Physiology of Cortical Neurons. Neuroscience 388, 23–35 (2018).

37. Wingert, J. C. & Sorg, B. A. Impact of Perineuronal Nets on Electrophysiology of Parvalbumin Interneurons, Principal Neurons, and Brain Oscillations: A Review. Front. Synaptic Neurosci. 13, 673210 (2021).

38. Giamanco, K. A., Morawski, M. & Matthews, R. T. Perineuronal net formation and structure in aggrecan knockout mice. Neuroscience 170, 1314–1327 (2010).

39. Lev-Ram, V. et al. Do Perineuronal Nets Stabilize the Engram of a Synaptic Circuit? Cells 13, 1627 (2024).

40. Berry, C. T., Sceniak, M. P., Zhou, L. & Sabo, S. L. Developmental Up-Regulation of Vesicular Glutamate Transporter-1 Promotes Neocortical Presynaptic Terminal Development. PLOS ONE 7, e50911 (2012).

41. Kawaguchi, Y., Katsumaru, H., Kosaka, T., Heizmann, C. W. & Hama, K. Fast spiking cells in rat hippocampus (CA1 region) contain the calcium-binding protein parvalbumin. Brain Research 416, 369–374 (1987).

42. Yizhar, O. et al. Neocortical excitation/inhibition balance in information processing and social dysfunction. Nature 477, 171–178 (2011).

43. Cao, W. et al. Gamma Oscillation Dysfunction in mPFC Leads to Social Deficits in Neuroligin 3 R451C Knockin Mice. Neuron 97, 1253–1260.e7 (2018).

44. Liu, L. et al. Cell type–differential modulation of prefrontal cortical GABAergic interneurons on low gamma rhythm and social interaction. Science Advances 6, eaay4073 (2020).

45. Hrvatin, S. et al. Single-Cell Analysis of Experience-Dependent Transcriptomic States in Mouse Visual Cortex. Nat Neurosci 21, 120–129 (2018).

46. Ye, L. et al. Wiring and Molecular Features of Prefrontal Ensembles Representing Distinct Experiences. Cell 165, 1776–1788 (2016).

47. Ueno, H. et al. Age-dependent and region-specific alteration of parvalbumin neurons and perineuronal nets in the mouse cerebral cortex. Neurochemistry International 112, 59–70 (2018).

48. Ueno, H., Suemitsu, S., Okamoto, M., Matsumoto, Y. & Ishihara, T. Parvalbumin neurons and perineuronal nets in the mouse prefrontal cortex. Neuroscience 343, 115–127 (2017).

49. Carulli, D. et al. Animals lacking link protein have attenuated perineuronal nets and persistent plasticity. Brain 133, 2331–2347 (2010).

50. Galtrey, C. M., Kwok, J. C. F., Carulli, D., Rhodes, K. E. & Fawcett, J. W. Distribution and synthesis of extracellular matrix proteoglycans, hyaluronan, link proteins and tenascin-R in the rat spinal cord. European Journal of Neuroscience 27, 1373–1390 (2008).

51. Brakebusch, C. et al. Brevican-Deficient Mice Display Impaired Hippocampal CA1 Long-Term Potentiation but Show No Obvious Deficits in Learning and Memory. Molecular and Cellular Biology 22, 7417–7427 (2002).

52. Favuzzi, E. et al. Activity-Dependent Gating of Parvalbumin Interneuron Function by the Perineuronal Net Protein Brevican. Neuron 95, 639–655.e10 (2017).

53. Sanchez, B., Kraszewski, P., Lee, S. & Cope, E. C. From molecules to behavior: Implications for perineuronal net remodeling in learning and memory. Journal of Neurochemistry 168, 1854–1876 (2024).

54. Dankovich, T. M. & Rizzoli, S. O. The Synaptic Extracellular Matrix: Long-Lived, Stable, and Still Remarkably Dynamic. Front. Synaptic Neurosci. 14, 854956 (2022).

55. Vigneault, É. et al. Distribution of vesicular glutamate transporters in the human brain. Front. Neuroanat. 9, (2015).

56. Herzog, E. et al. In Vivo Imaging of Intersynaptic Vesicle Exchange Using VGLUT1Venus Knock-In Mice. J Neurosci 31, 15544–15559 (2011).

57. Khan, M., Sevilla, L. D., Mahesh, V. B. & Brann, D. W. Enhanced Glutamatergic and Decreased Gabaergic Synaptic Appositions to GnRH Neurons on Proestrus in the Rat: Modulatory Effect of Aging. PLOS ONE 5, e10172 (2010).

58. Misselhorn, J., Schwab, B. C., Schneider, T. R. & Engel, A. K. Synchronization of Sensory Gamma Oscillations Promotes Multisensory Communication. eNeuro 6, ENEURO.0101-19.2019 (2019).

59. Sedley, W. & Cunningham, M. O. Do cortical gamma oscillations promote or suppress perception? An under-asked question with an over-assumed answer. Front. Hum. Neurosci. 7, (2013).

60. Anand, S. et al. High gamma coherence between task-responsive sensory-motor cortical regions in a motor reaction-time task. Journal of Neurophysiology 130, 628–639 (2023).

61. Gupta, D. S. & Chen, L. Brain oscillations in perception, timing and action. Current Opinion in Behavioral Sciences 8, 161–166 (2016).

62. Başar, E., Başar-Eroglu, C., Karakaş, S. & Schürmann, M. Gamma, alpha, delta, and theta oscillations govern cognitive processes. International Journal of Psychophysiology 39, 241–248 (2001).

63. Donato, F., Rompani, S. B. & Caroni, P. Parvalbumin-expressing basket-cell network plasticity induced by experience regulates adult learning. Nature 504, 272–276 (2013).

64. Aouci, R. et al. The Antidepressant Action of Fluoxetine Involves the Inhibition of Dlx5/6 in Cortical GABAergic Neurons through a TrkB-Dependent Pathway. Cells 13, 1262 (2024).

65. Lee, P. R. & Fields, R. D. Activity-Dependent Gene Expression in Neurons. Neuroscientist 27, 355–366 (2021).

66. Benes, F. M. & Berretta, S. GABAergic interneurons: implications for understanding schizophrenia and bipolar disorder. Neuropsychopharmacology 25, 1–27 (2001).

67. Marín, O. Interneuron dysfunction in psychiatric disorders. Nat Rev Neurosci 13, 107–120 (2012).

68. Ichim, A. M. et al. The gamma rhythm as a guardian of brain health. eLife 13, e100238 (2024).

69. Benes, F. M., McSparren, J., Bird, E. D., SanGiovanni, J. P. & Vincent, S. L. Deficits in small interneurons in prefrontal and cingulate cortices of schizophrenic and schizoaffective patients. Arch Gen Psychiatry 48, 996–1001 (1991).

70. Pantazopoulos, H., Woo, T.-U. W., Lim, M. P., Lange, N. & Berretta, S. Extracellular matrix-glial abnormalities in the amygdala and entorhinal cortex of subjects diagnosed with schizophrenia. Arch Gen Psychiatry 67, 155–166 (2010).

71. Pantazopoulos, H. et al. Aggrecan and chondroitin-6-sulfate abnormalities in schizophrenia and bipolar disorder: a postmortem study on the amygdala. Transl Psychiatry 5, e496 (2015).

72. Hashemi, E., Ariza, J., Rogers, H., Noctor, S. C. & Martínez-Cerdeño, V. The Number of Parvalbumin-Expressing Interneurons Is Decreased in the Prefrontal Cortex in Autism. Cereb Cortex 27, 1931–1943 (2017).

73. Uhlhaas, P. J. & Singer, W. Abnormal neural oscillations and synchrony in schizophrenia. Nat Rev Neurosci 11, 100–113 (2010).

74. Barr, M. S. et al. Evidence for excessive frontal evoked gamma oscillatory activity in schizophrenia during working memory. Schizophr Res 121, 146–152 (2010).

75. Haenschel, C. et al. Cortical oscillatory activity is critical for working memory as revealed by deficits in early-onset schizophrenia. J Neurosci 29, 9481–9489 (2009).

