## Supplementary Figures for "*Dlx5/6* regulate perineuronal net-synapse coupling and stabilize adult cortical Parvalbumin neurons networks"

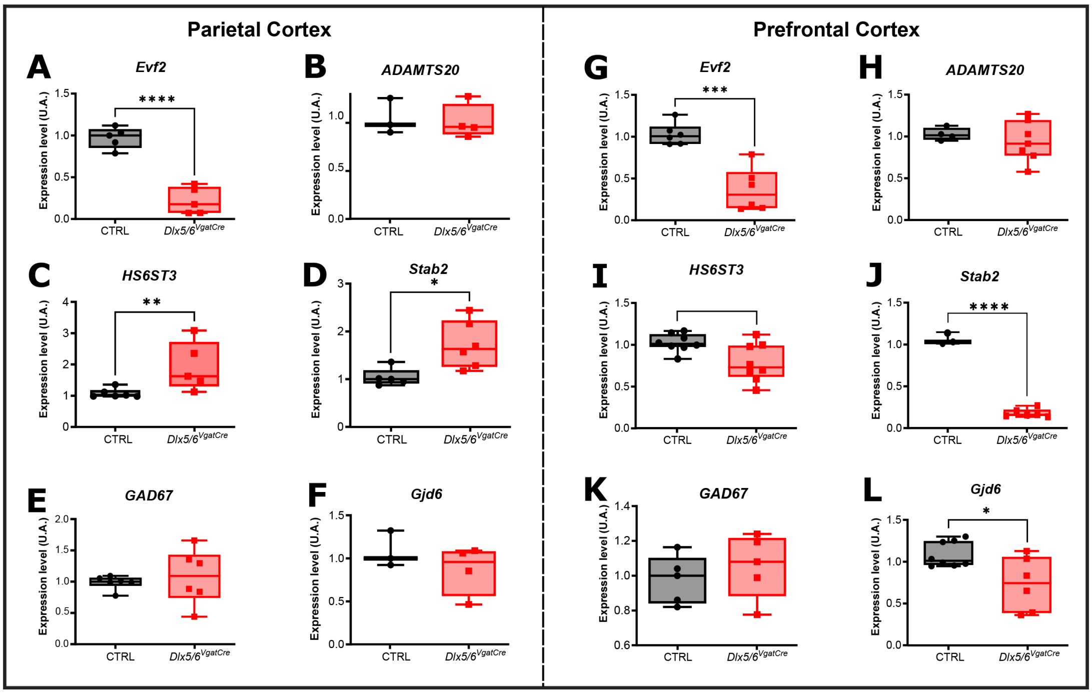


**Supp Figure 1.** ***Dlx5/6*** **inactivation differentially regulates the expression of extracellular matrix- and GABAergic-related genes in the somatosensory and prelimbic cortices.**
(**A-F**) RT-qPCR analysis of relative gene expression in the somatosensory cortex (S1BF) of control (CTRL) and *Dlx5/6*$^{VgatCre}$ mice for *Evf2b* (**A**), *Adamts20* (**B**), *Hs6st3* (**C**), *Stab2* (**D**), *Gad1* (**E**), and *Gjd6* (Connexin 30) (**F**). (**G-L**) RT-qPCR analysis of relative gene expression in the prelimbic cortex (PrL) of CTRL and *Dlx5/6*$^{VgatCre}$ mice for *Evf2* (**G**), *Adamts20* (**H**), *Hs6st3* (**I**), *Stab2* (**J**), *Gad1* (**K**), and *Gjd6* (Connexin 30) (L). Data are shown as box-and-whisker plots, with the box indicating the median and interquartile range, and whiskers extending to the minimum and maximum values (n = 6 mice per group). Statistical significance was assessed using unpaired two-tailed Student's t-tests. *p < 0.05, **p < 0.01, ***p < 0.001, ****p < 0.0001.

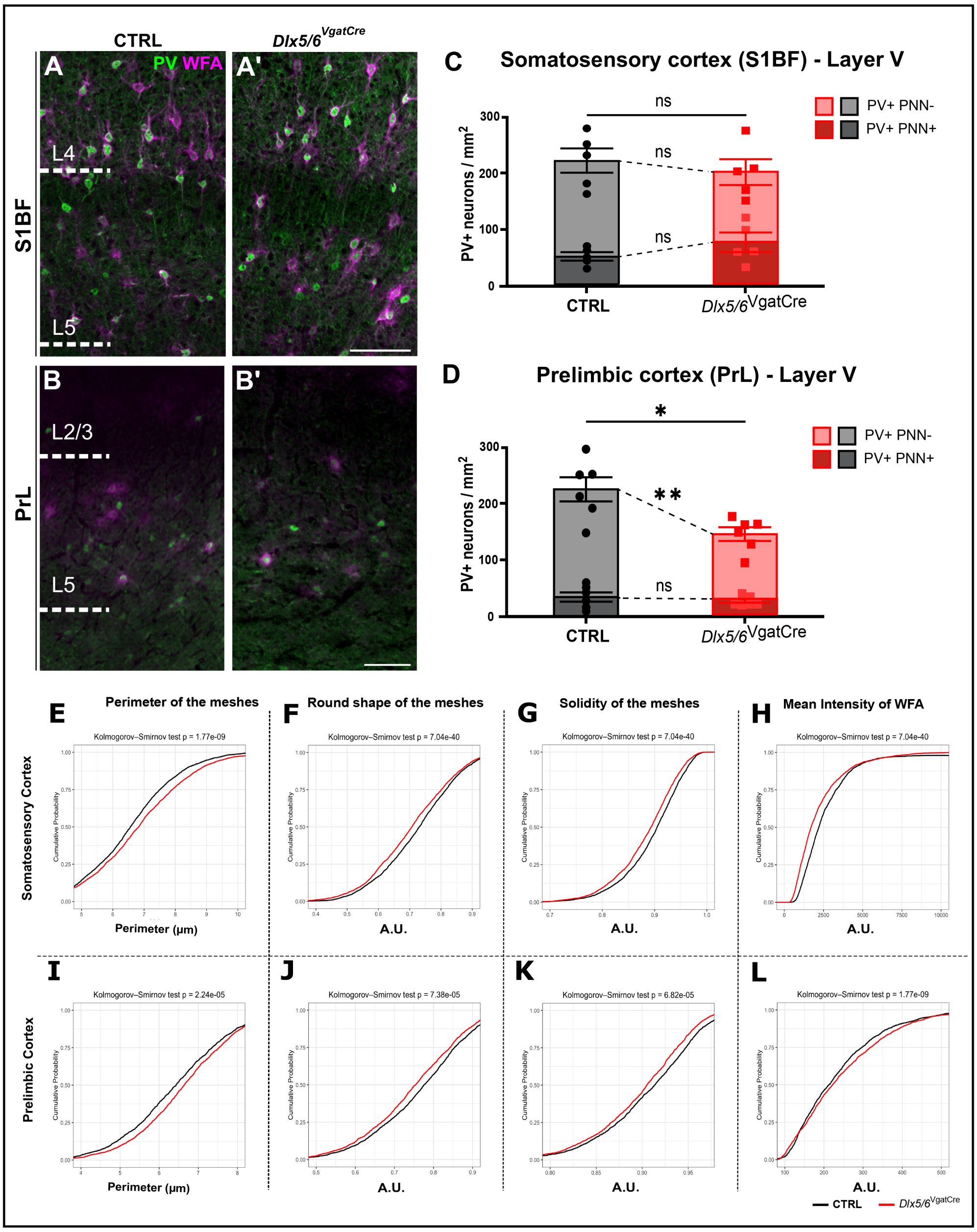


**Supp Figure 2. Region specific *Dlx5/6* inactivation consequences on PV-neurons and PNN$^{+}$ neuron density, and shifts the cumulative distributions and intensity of PNN mesh in both cortical regions.** (**A-B'**) Representative double immunofluorescence images of Parvalbumin-positive (PV$^{+}$; green) interneurons and perineuronal nets (PNN$^{+}$; WFA; magenta) in the primary somatosensory barrel field cortex (S1BF; **A-A'**) and prelimbic cortex (PrL; **B-B'**) of control and *Dlx5/6$^{VgatCre}$* mice. Dashed lines indicate cortical layer boundaries. (**C**) Quantification of PV$^{+}$PNN$^{-}$ and PV$^{+}$PNN$^{+}$ cell densities in the S1BF revealed no significant differences between genotypes. (**D**) In the prelimbic cortex, *Dlx5/6* inactivation significantly reduced total PV$^{+}$ cell density, driven by a decrease in PV$^{+}$PNN$^{-}$ cells, whereas PV$^{+}$PNN$^{+}$ cell density remained unchanged relative to controls. (**E-H**) Cumulative distribution analysis of PNN mesh parameters and WFA fluorescence intensity in the S1BF of control and *Dlx5/6$^{VgatCre}$* mice. *Dlx5/6* inactivation was associated with a rightward shift in mesh perimeter distribution, indicating larger mesh size (**E**), alongside alterations in mesh roundness (**F**) and solidity (**G**). WFA intensity distribution was shifted leftward, indicating reduced labelling intensity within PNN meshes (**H**). (**I-L**) In the PrL, mesh perimeter (**I**), roundness (**J**), and solidity (**K**) distributions were similarly altered between genotypes, together with a leftward shift in WFA intensity (**L**). Data are presented as mean ± SEM (n = 5 mice per group). PV$^{+}$ cell density differences were assessed using unpaired two-tailed Student's t-tests; cumulative distribution differences were assessed using two-sample Kolmogorov–Smirnov tests. *$p < 0.05$; **$p < 0.01$; n.s., not significant. Scale bars, 50 μm.

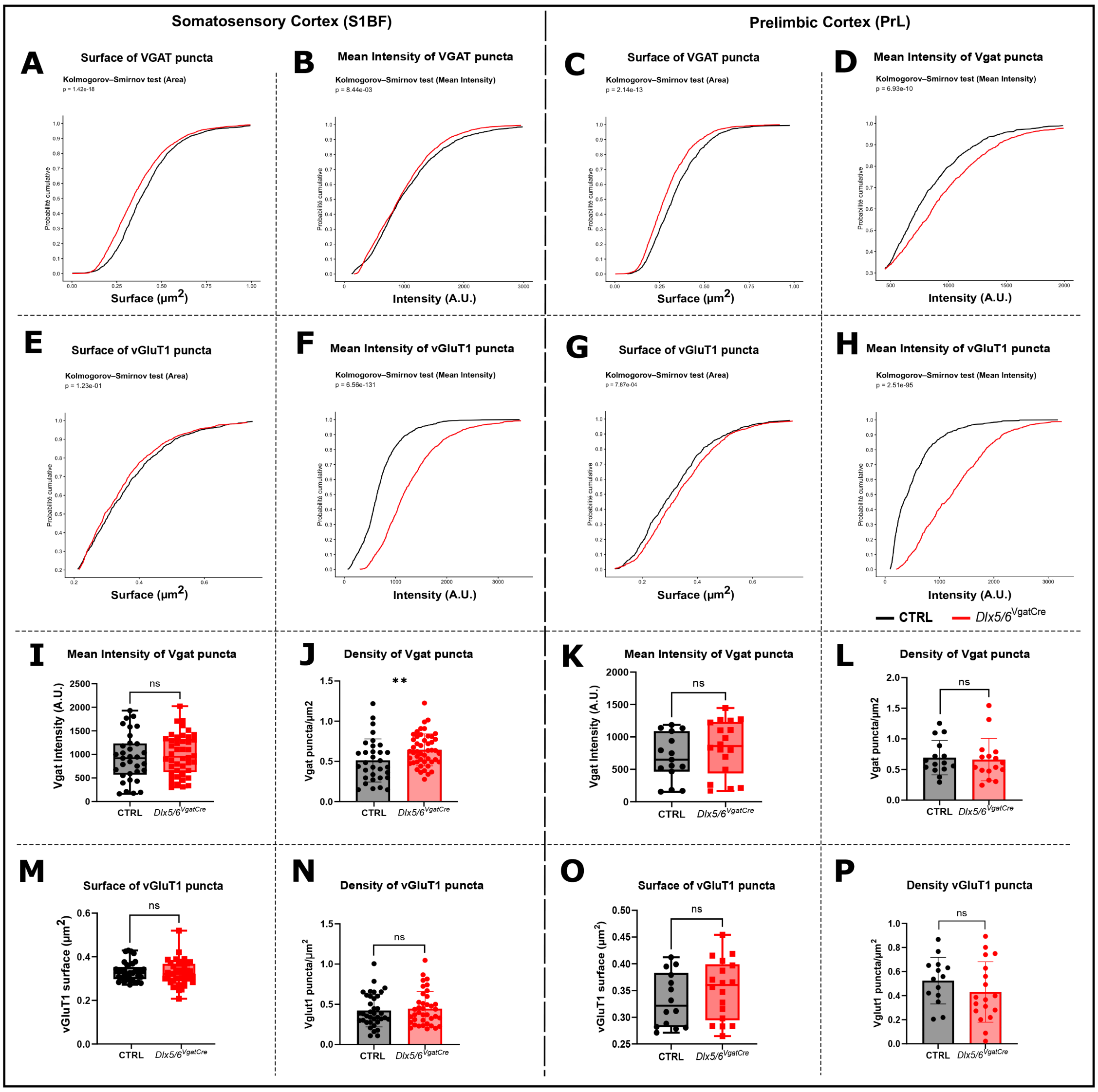


**Supp Figure 3. Detailed characterization of inhibitory VGAT and excitatory VGLUT1 presynaptic puncta parameters in the S1BF and PrL cortices.**

**(A-D)** Cumulative distribution analysis of VGAT puncta surface area (**A, C**) and mean fluorescence intensity (**B, D**) in the S1BF (**A, B**) and PrL (**C, D**) of control and *Dlx5/6$^{VgatCre}$* mice. (E–H) Cumulative distribution analysis of VGLUT1 puncta surface area (**E, G**) and mean fluorescence intensity (**F, H**) in the S1BF (**E, F**) and PrL (**G, H**). (**I-L**) Mean VGAT puncta intensity (**I, K**) and puncta density (**J, L**) per cell in the S1BF (**I, J**) and PrL (**K, L**). (**M-P**) VGLUT1 puncta surface area (**M, O**) and puncta density (**N, P**) per cell in the S1BF (**M, N**) and PrL (**O, P**). For (**A-H**), statistical significance was assessed using two-sample Kolmogorov–Smirnov tests; exact p-values are indicated on each panel. For (**I-P**), Permutation test using Monte Carlo method (9,999 permutations) were used. Data are presented as mean ± SEM. $^{**}p < 0.01$; ns, not significant.

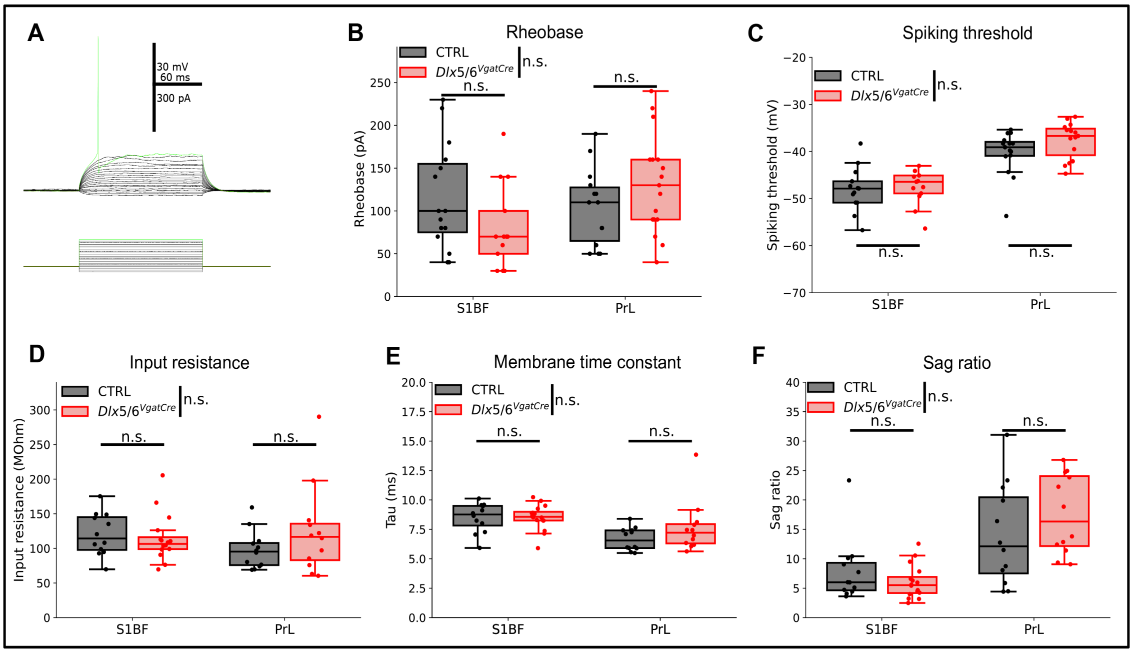


**Supp Figure 4. *Dlx5/6* inactivation in VGAT$^{+}$ neurons leave fast-spiking interneurons (FS-INs) spiking threshold and passive properties unaffected.**

**A**) An example recording for finding the rheobase (first current intensity evoking an action potential). **B**) Rheobase quantification. 2-way ANOVA reveals no effect of the cortex or the genotype, but reveals a significant interaction between genotype and cortex (p=0.034075). However, within cortex comparison (T-tests) does not reveal any difference between the two genotypes. **C)** Spiking threshold quantification. 2-way ANOVA reveals no genotype effect (p=0.15) nor interaction between genotype and cortex (p=0.36). Within cortex comparison (PrL: permutation tests; S1BF: T-test) reveals no significant differences between genotypes. **D**) Input resistance quantification. 2-way ANOVA reveals no genotype effect (p=0.41) nor interaction between genotype and cortex (p=0.16). Within cortex comparison (T-tests) reveals no significant differences between genotypes. **E**) Membrane time constant quantification. 2-way ANOVA reveals no genotype effect (p=0.23) nor interaction between genotype and cortex (p=0.24). Within cortex comparison (t-tests) reveals no significant differences between genotypes. **F**) Sag ratio quantification. 2-way ANOVA reveals no genotype effect (p=0.71) nor interaction between genotype and cortex (p=0.12). Within cortex comparison (t-tests) reveals no significant differences between genotypes. ns, not significant.

**Supplementary Table 1.** Primers used in this study for RT-qPCR.

| | |
|---|---|
| *Acan* | Fw: 5’ TGG ATC GGT CTG AAT GAC AGG 3’<br>Rv: 5’ AGA AGT TGT CAG GCT GGT TTG G 3’ |
| *Bcan* | Fw: 5’ GAG GAT CGA GCC TTC CGC 3’<br>Rv: 5’ GGT GGA CGT GGC ATG GGA T 3’ |
| *TnR* | Fw: 5’ CAC TCC AAA GAA CAA TGA AG 3’<br>Rv: 5’ GCT TGT TCC TTT AGT GCT AC 3’ |
| *HAPLN* | Fw: 5’ CTG GAG GAT TAT GGA AGA TA 3’<br>Rv: 5’ CAC CAC ACC TTG TAA CTC TA 3’ |
| *Has3* | Fw: 5’ CCT TGG CAA CTC AGT GGA CTA C 3’<br>Rv: 5’ TGG ACA TCT CCT CCA ACA CCT C 3’ |
| *MMP9* | Fw: 5’ GCT CCT GGC TCT CCT GGC TT 3’<br>Rv: 5’ GTC CCA CCT GAG GCC TTT GA 3’ |
| *PV* | Fw: 5’ TGT CGA TGA CAG ACG TGC TC 3’<br>Rv: 5’ TTC TTC AAC CCC AAT CTT GC 3’ |
| *Evf2B* | Fw: 5’ CTC CCT CCG CTC AGT ATA GAT TTC 3’<br>Rv: 5’ CCT CCC CGG TGA ATA TCT CTT 3’ |
| *ADAMST20* | Fw: 5’ GTC CTG GGA AGT TCG TTT CCA 3’<br>Rv: 5’ GGC TGA AAT GCC GGT TCT G 3’ |
| *ADAMTS11* | Fw: 5’ GCC CAG GTG AGT TTC GTC ATC 3’<br>Rv: 5’ GCT CCA CAT ACT GCG AGG A 3’ |
| *HS6ST3* | Fw: 5’ CCA TCA TGG AGA AGA AGG AT 3’<br>Rv: 5’ GTA GGC AGC TCA TCT GGT GT 3’ |
| *Stab2* | Fw: 5’ AGC TGC TGC CTT TAA TCC TCA 3’<br>Rv: 5’ ACT CCG TCT TGA TGG TTA GAG TA 3’ |
| *GAD67* | Fw: 5’ TGG CAT CTT CCA CTC CTT CG 3’<br>Rv: 5’ GGC TAC GCC ACA CCA AGT AT 3’ |
| *Cnx30* | Fw: 5’ ACCAGCATAGGGAAGGTGTG 3’<br>Rv: 5’ TGC AGA GTG TTG CAG ACA AAG 3’ |
| *Actin B* | Fw: 5’ CAT TGC TGA CAG GAT GCAGAAGG 3’<br>Rv: 5’ TGC TGG AAG GTG GAC AGT GAG G 3’ |
| *ß3-Tubulin* | Fw: 5’ CAT CAG CGA TGA GCA CGG CAT A 3’<br>Rv: 5’ GGT TCC AAG TCC ACC AGA ATG G 3’ |
